# Biological embedding of childhood adversity - a multi-omics perspective on stress regulation

**DOI:** 10.1101/2023.06.10.544462

**Authors:** Johannes C.S. Zang, Caroline May, Katrin Marcus, Robert Kumsta

**Affiliations:** Faculty of Psychology, Institute for Health and Development, Ruhr University Bochum, Bochum, Germany; Department of Behavioural and Cognitive Sciences, Laboratory for Stress and Gene-Environment Interplay, University of Luxemburg, Esch-sur-Alzette, Luxemburg; DZPG (German Center for Mental Health), partner site Bochum/Marburg, Germany; Medizinisches Proteom-Center, Medical Proteome Analysis Centre for Protein Diagnostics (PRODI), Ruhr University Bochum, Germany

**Author notes:** Corresponding address: Prof Dr. Robert Kumsta, Faculty of Humanities, Education and Social Sciences, Maison des Sciences Humaines, 11, Porte des Sciences, L-4366 Esch-sur-Alzette.

**Keywords:** Multi-omics, childhood adversity, stress exposure, monocytes, transcriptomics, proteomics, DNA methylation

## Abstract

The experience of adversity in childhood can have life-long consequences on health outcomes. In search of mediators of this relationship, alterations of bio-behavioral and cellular regulatory systems came into focus, including those dealing with basic gene regulatory processes. Systems biology oriented approaches have been proposed to gain a more comprehensive understanding of the complex multiple interrelations between and within layers of analysis. We used co-expression based, supervised and unsupervised single and multi-omics system approaches to investigate the influence of childhood adversity on gene expression, protein expression and DNA methylation in CD14^+^ monocytes of healthy adults before and after exposure to an experimental psychosocial stress protocol. Childhood adversity explained some variance at the single analyte level and within gene and protein co-expression structures. A single-omic, post stress gene expression model differentiated best between participants with a history of childhood adversity and controls in supervised analyses. In unsupervised analyses, a multi-omics based model showed best performance but separated participants based on sex only. Multi-omics analyses are a promising concept but might yield different results based on the specific approach taken and the omic-datasets supplied. Here, stress associated gene-expression pattern were most strongly associated with childhood adversity, and integrating multiple cellular layers did not results in better discriminatory performance. Currently, the capacity and yield of different omics-profiling methods might limit the full potential of integrative approaches.

## Introduction

Exposure to stressful and potentially traumatic events in childhood not only causes immediate harm but also represents a trans-diagnostic risk factor for a range of mental health problems across the life span (1, 2). Furthermore, early adversity and trauma is among most potent and consistent predictors of physical disease and reduced longevity (3). These long-term effects and also the stability of early environmental insults has led to the question of, how and when the social becomes biological ‘; or in other words, how psychosocial risk becomes *biologically embedded*. There is evidence that that early adversity during different periods of developmental plasticity induces enduring changes at multiple levels of regulatory systems, and influences social, cognitive, neuronal and emotional development (4).

Among the bio-behavioural systems involved in bi-directional mind-body interplay, alterations of stress response systems, inflammatory immune processes and the stress-immune interface have emerged as likely mediators of the effects of adverse childhood experiences on health outcomes (5).

More recently, the sub-cellular realm came into focus. Advances in research on animal models and technological developments - especially high throughput -*omics* approaches provided a framework to investigate the immediate and fundamental mechanisms of gene-environment interplay. Particularly epigenetic modification such as DNA methylation received attention (6) and RNA dynamics are increasingly being studied (7).

*Omics*-technology driven access to the entirety of the biochemical landscape of different molecular layers within cells, such as the transcriptome or proteome (8), also fuel the realization of system biology approaches in the study of complex traits or environmental influence (9).

System biology approaches emphasize on describing the interrelation of analytes or cell entities to gain insight into the complexity of biological processes underlying phenotypic variation and disease risk (10). A steadily increasing number of statistical methods allows for the implementation of these perspectives in research targeting one or more than one molecular layers (11).

Co-expression network analyses for example reflect a systems approach to single - *omics* analyses and aim to identify network structures formed by highly interrelated analytes. As such, weighted gene-co-expression network analysis (WGCNA) has been used to gain insight into gene co-expression signatures associated with posttraumatic stress disorder (PTSD) (12).

Multi-*omics* analyses further advance the idea of molecular systems biology. Such approaches integrate three or more (11) -*omics* datasets derived from specific profiling methods, for example the application of microarrays or mass spectrometry. Although the integration of several large datasets with different modalities impose a certain kind of complexity to the analytical enterprise (9, 13), it promises a more comprehensive insight into the molecular machinery unmet by a single-omics based perspective.

More broadly, such multi-*omic* analyses can be classified into supervised and unsupervised approaches (11, 14).

Unsupervised approaches are exploratory and data driven. They do not require precise specification of the clinical/experimental group from which the included omics datasets are derived, but may be used e.g. to reveal clusters of participants within the sample. The application of regularized unsupervised multiple kernel learning (rMKL) to gene expression, DNA methylation and miRNA data for example resulted in the identification of sub-groups in patients with distinct cancer types, which showed different survival times (15).

Supervised multi-omics approaches, on the other hand, are hypothesis-driven and seek to identify molecular signatures that best characterize a known and specified clinical/experimental group. Thus, a supervised strategy is particularly suited to identify predictive biomarkers or condition specific molecular processes. *D*ata *i*ntegration *a*nalysis for *b*iomarker discovery using *l*atent c*o*mponents (DIABLO; 16), for example represents a specific supervised approach which has been used to identify molecular signatures predictive of Hepatitis B vaccination response (17).

Taken together, despite increasing insights into adversity-related biological alterations, knowledge is fragmentary and mostly based on single levels of analysis. Integrative approaches grounded in a systems biology perspective promise to enhance our understanding of alterations within gene and cell regulatory mechanisms following experience to childhood adversity Here, we apply such integrative approaches to childhood adversity in the context of acute stress exposure, taking into account stress-associated hormonal response dynamic and the associated immediate changes on the level of transcriptome and proteome. Based on cortisol measures, DNA methylation, gene expression and protein expression data derived from CD 14^+^ monocytes, we showcase the application of i) single-omics co-expression network analysis; ii) unsupervised and iii) supervised multi-omics analysis to identify acute stress-associated molecular signatures in adults reporting childhood adversity

## Methods

### Study design and context

This study is based on a project investigating long-term consequences of childhood adversity on stress reactivity, which included the collection of transcriptomic, proteomic, and genomewide DNA methylation data. Initial work focused on individual omics data (see supplements; (18, 19, 20). Here we take integrative multi-omics approaches to further characterize molecular signatures associated with adverse childhood and acute stress experience. The analyses reported here incorporate gene and protein expression data derived from monocytes collected during a baseline measurement minutes 45 min. before (t0) and 180 min. following exposure to psychosocial stress (t1). Genome-wide DNA methylation data derives from monocytes collected 45 min after stress exposure. Cortisol measures were taken before and after exposure to the Trier Social Stress Test (TSST), at -45, -2, 1, 10, 20, 30, 45, and 90 min relative to stress exposure (Figure 1A).

**Figure 1:**
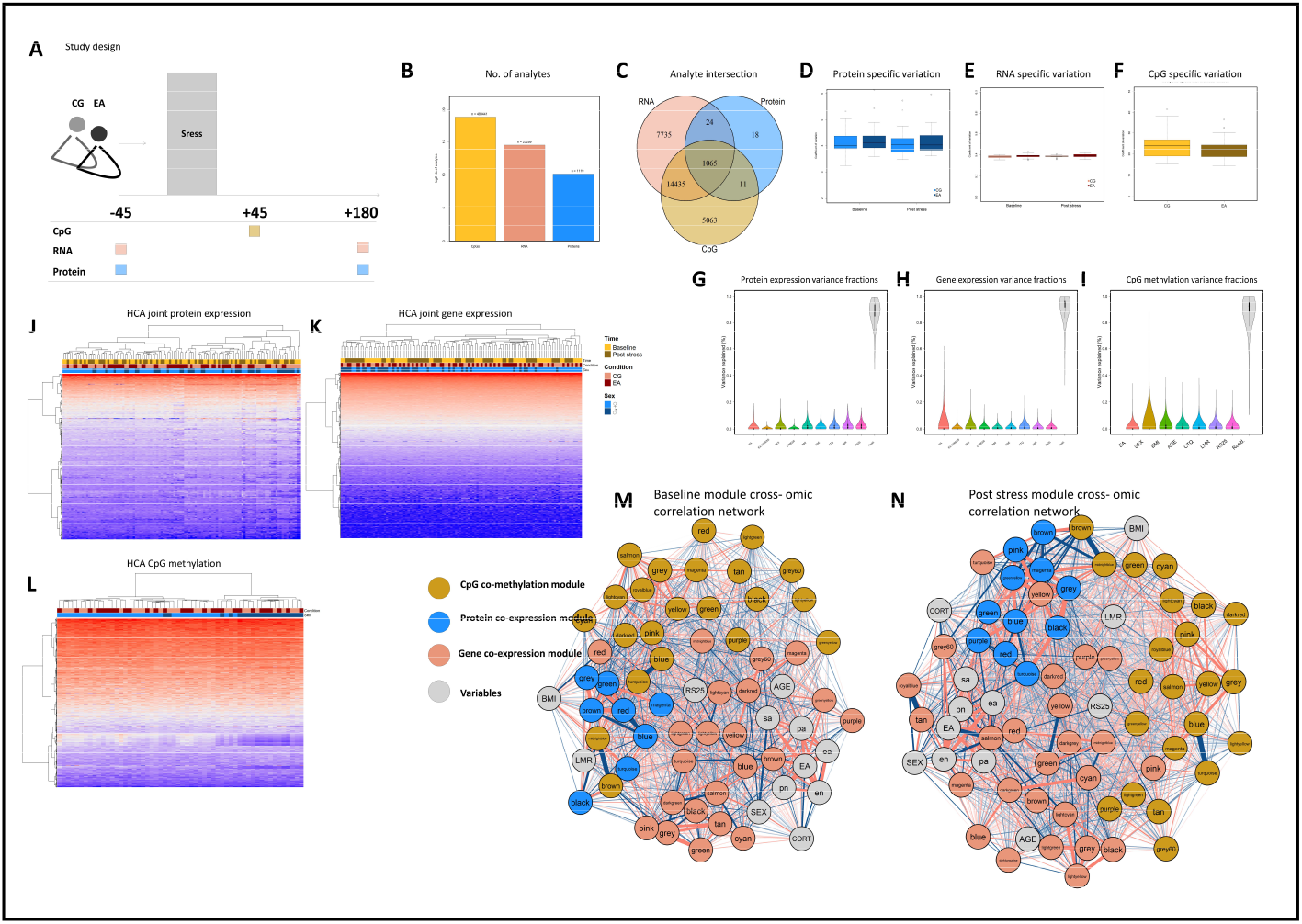
The study design depicts major sampling points of monocytes for transcriptome (RNA = red), proteome (protein = blue) and whole genome CpG methylation (CpG = yellow) analysis in minutes relative to stress exposure (TSST) (A). The number of analytes and intersection of gene names assigned to analytes across omic-datasets is given in (C). Boxplots depict condition specific inter-individual variation (CV) in baseline and post stress gene & protein expression (C & E) and variation in CpG methylation (G). Variance fractions explained by clinical variables within protein (D) and gene (F) expression data and CpG methylation (H) is presented by violin plots. Networks show baseline (I) and post stress (J) correlation (edges) of proteome (blue), transcriptome (red) or CpG methylation (yellow) derived co-expression (co-methylation) modules (nodes). Grey nodes represent variables of interest (see methods).

### Sample

The sample consisted of 60 healthy adults reporting a history of childhood adversity (EA) and a matched control group (CG). Self-reported childhood adversity was assessed with the Childhood Trauma Questionnaire (CTQ) followed by a more in-depth interview using the Early Trauma Inventory (ETI). All participants were screened for mental disorders using Structured Clinical Interview for DSM Disorders (SKID I & II) and filled in the Resilience Scale (RS-25). Recruiting and sampling procedures as well as the assessment of further bio-behavioral variables are described in more detail elsewhere (18). See supplements for sample characteristics.

### Omics profiling

DNA, mRNA and proteins were extracted from CD14+ monocytes, isolated from blood samples via immunomagnetic cell separation. DNA methylation levels were quantified with the llumina Infinium HumanMethylation450 BeadChip and processed with standard bioinformatic pipelines. Following RNA isolation, genome-wide gene expression profiling was realized using Agilent Whole Human Genome Oligo Microarrays 8x60K V2. Protein was extracted from monocytes using repeated exposure to ultrasonic waves and the application of physical force under precooled and iced conditions. Proteome analysis was realized by Liquid Chromatography tandem Mass spectrometry (see(20) and supplements for details).

### Final datasets

Following preprocessing, we constructed the final sample as intersection of participants with valid values on all omics datasets (N = 56) and mapped individual analytes (CpG sites, transcripts, proteins) to gene symbols were possible. Additional variables included group assignment (early adversity vs control), CTQ scores, RS25 scores, BMI, age, sex, *monocyte*/*lymphocyte ratio* (MLR) and time of assessment (t0/t1) relative to stress. Participants endocrine stress reaction was coded as cortisol increase following TSST exposure (baseline-to-peak levels).

### Analytical strategy

We first assessed group specific inter-individual variance and analyte specific sources of variance. We approached integration of our transcriptomic, proteomic, and CpG methylation data in three ways. First, we utilized co-expression network analysis to identify modules of co-expressed analytes on the single omics level and subsequently investigated the cross-omics interrelation of level specific co-expression structures. Second, following an unsupervised pathway to multi-omics integration, we utilized regularized multiple kernel learning for dimensionality reduction (rMKL-DR) and tested how good rMKL-DR differentiates between participants when supplied with gene expression, protein expression, CpG methylation and cortisol response data. Third, following a supervised pathway to multi-omics integration, we utilized sparse and partial least squares regression discriminant analysis to identify single- or multi-omics derived subsets of analytes that discriminate best between participants with a history of childhood adversity and control participants.

### Analysis of stress related and condition-specific interpersonal variation

As a measure of interpersonal variation, coefficients of variation (CV) were calculated as CV

= standard deviation/mean on non-transformed raw datasets. Proteome and gene expression specific CV were calculated on globally normalized gene (quantile) and protein expression (LFQ) data, separately for baseline and post stress measurements. DNA methylation specific CV were calculated on basis of m-values. For each analyte and point of measurement CV were determined for participants with a history of childhood adversity (EA) and control participants (CG) respectively. We calculated Pearson’s r to investigate the interrelation between the analyte and time specific CV magnitude and compared variation in baseline and post stress analyte expression between groups using asymptotic testing as implemented in the cvequality R package (version 0.1.3; (21)).

### Omic specific variance analysis

We used a multiple regression model approach to delineate sources of variation and specifically characterize the extent of variance in CpG methylation, gene expression or protein expression attributable to variables of interest. As such, all applied models included participants’ age, sex, BMI, LMR, RS25 scores and group (EA/CG). Two versions of these models were specified, including either participants CTQ total score or participants’ scores on the CTQ subdimensions. Models designed for the analysis of stress effects additionally accounted for the time of measurement and were extended to include the variable stress (t0/t1) as well as an intersection effect (stress: condition). In all models, sex, condition, stress and the stress:condition interaction were specified as random effects and models were fitted using the R package variancePartition (version 1.22.0; (22)).

### Co-expression and co-methylation network analysis

Co-expression and co-methylation network analyses were conducted to examine correlation patterns in the respective omics datasets using weighted correlation network analysis by means of the WGCNA R package (version 1.70-3; (23)). Module eigengenes, eigenproteins and eigenCpGs (MEs) were calculated for each module and used to identify modules related to the experience of early adversity or relevant other variables (modules of interest) and to investigate cross-omics interrelation of identified co-expression modules. We used HCA to identify clusters of correlated modules and constructed two undirected correlation networks using the R qgraph package (version 1.9; (24)) to visualize cross-omics module interrelation and module – variable interrelations in baseline or post stress gene and protein co-expression modules and co-methylation modules (see supplements for details).

### An unsupervised pathway to integration. The regularized multiple kernel learning for dimensionality reduction analysis approach (rMKL-DR)

rMKL-DR was performed using Locality Preserving Projections (LPP) for dimensionality reduction as suggested by Speicher and Pfeifer (15). All analyses were conducted within four different clusters, containing different omic levels. The first cluster (M1 - M3) contained protein expression and gene expression data. The second cluster adds participants’ base-to-peak cortisol response to protein expression and gene expression data (M4 - M6). The third cluster included gene expression, protein expression, and methylation data (M7 - M9). The fourth cluster (M10 - M12) integrated all data sets. We run three analyses for each cluster with either two, three or five (default setting) projection dimension and all analyses were performed separately for baseline and post stress data. Across all analyses, the number of neighbours was hold at a constant of nine (default setting). We used multiple regression modelling (see above) to delineate sources of variation of rMKL derived subgroups (see supplements for details).

### A supervised pathway to integration: The Data Integration Analysis for Biomarker discovery using Latent variable approaches for Omics studies (DIABLO) approach

We utilized supervised discriminant analyses as implemented in the R mixOmics package (version 6.16.3; (25)) to identify analytes explaining participants group membership (EA/CG). Analyses were performed either on each of the omics datasets separately (R = Gene expression data, P = protein expression data, M = Methylation data) using sparse partial least squares regression discriminant analysis (sPLSDA), on gene and protein expression data (RP) or gene expression, protein expression and CpG methylation data (RPM) simultaneously using block sparse partial least squares regression discriminant analysis (block sPLSDA) as provided by the DIABLO framework (16). All analyses were performed separately on baseline and post stress data. To select the model with the best discriminant performance, we run each model ten times while increasing the number of selected analytes and evaluated the model performance on basis of balanced error rate (BER, smaller = better).

## Results

Monocyte-derived multi-omics datasets comprised altogether 504.819 analytes including expression values of 23.259 transcripts, 1.119 proteins, and methylation degrees of 480.441 CpGs. 1.065 of these analytes mapped to the same gene name (Figure 1B).

### Omic specific inter-individual variability

The assessment of inter-individual variability (CV) across all omics datasets showed highest amount of variation within CpG methylation degrees and lowest variation within gene expression levels (Figure 1C,E,G). Asymptotic tests did not reveal any significant group differences in expression or DNA methylation related variation (Table S4) and there was no significant relation between inter-individual and time specific variability across omics datasets in participants with a history of childhood adversity. In control individuals, variation in post stress protein expression was significantly correlated with variation in baseline protein expression (*r* = 0.73, *p*< .001) and baseline gene expression (*r* = -0.39, *p*< .05) (Table S2-3).

### Omic specific sources of variance

After controlling for all other variables included in the respective models, early adversity explained a modest fraction of variance in expression levels (gene, protein) or DNA methylation levels (Table S5-9). The same was true for the effects of stress exposure and the early adversity by stress interaction (Table S8 & 9).

### Analyte specific co-expression and co-methylation structures and cross-omic interrelation

WGCNA led to the identification of between 7 and 22 modules harboring co-expressed genes (n_t0_ = 21, n_t1_ = 22), co-expressed proteins (n_t0_ = 7, n_t1_ = 24) or co-methylated CpGs (n = 21) within the calculated networks (Figure S1 & 2). Uncharacterized analytes were assigned to a grey default module. Gene and protein co-expression structures identified in baseline networks were largely conserved within the post stress networks (Gene co-expression: Zsummary *M* = 20.39, *SD* = 11.83; Protein co-expression: Zsummary *M* = 11.06, *SD* = 4.43) and WGCNA conducted upon joint datasets revealed 21 gene co-expression modules and 6 protein co-expression modules.

Overall, both within and across layers, identified co-expression and co-methylation modules correlated moderately; however, strong correlations were observed for individual modules (range in baseline network: r= -0.93 to 0.93; post stress network: r= -0.92 to 0.86; joint network: r= -0,96 to 0.83). HCA driven analysis of module correlation indicated stronger within-omics than cross-omics relations (Figure S6). Interestingly, cluster structures showed a negative correlation between most of the co-methylation modules and particular gene and protein co-expression modules. This relation between higher DNA methylation degrees and reduced gene and protein expression became especially evident in baseline and joint data modules. Network structures derived from baseline (Figure 1I) and post stress modules (Figure 1J) validated these observations. Although modules of the same omics origin tended to cluster together within the topology of both pre- and post-stress networks, modules were stronger intertwined in the post stress network – indicated by higher strength and closeness and higher absolute edge weights (*t*(4497)= 11.46, *p* < .001) compared to the baseline network. In both networks, gene co-expression modules showed closest proximity to childhood adversity associated variables (EA or CTQ scores). A closer look at modules’ correlation with early adversity revealed the following: First, correlational analyses identified four baseline co-expression modules (two protein modules) positively associated with the experience of early adversity and analytes contained within these modules are functionally implicated in e.g. DNA strand elongation, interleukin-1 beta signaling viral processes and mitochondrial ATP synthesis. Second, following stress there was a larger number of gene (n = 5) and protein (n = 3) co-expression modules significantly correlated with a history of childhood adversity (Figure S4). Modules of interest harbored analytes involved in e.g. regulation of cellular response to stress, mitochondrial protein processing, viral response or neutrophil activation involved in immune response (Figure S7 & 8). Third, none of the identified co-methylation modules was associated with the experience of childhood adversity (Figure S4 A) and we found no relation between protein co-expression and acute stress exposure (Figure S5). For further details, see SI.

### An unsupervised pathway to integration. The regularized multiple kernel learning for dimensionality reduction analysis approach

Application of defined rMKL-DR models grouped participants in clusters of n = 2-10, based on baseline and post-stress omics data and participants’ cortisol response as input (Table 1). The groups formed by rMKL-DR tended to be larger when based on post-stress omics data. Table 1 shows the extent of the variance explained in the respective rMKL-DR cluster by variables of interest using multiple regression modelling. None of the models clearly differentiated between participants with a history of childhood adversity and the control group. The largest amount of variance in participants’ history of childhood adversity was explained by model 1 (6.14 %) based on RNA and Protein expression. Notably, including CpG methylation as data source clearly increased discriminatory capabilities of models particularly when it comes to differentiating participants based on the experience of sexual abuse, sex, age, or BMI. For instance, group assignments derived from constructed rMKL-DR models explained a substantial amount of variance of CTQ sexual abuse subscale scores (M7 = 29.3 %) and participants sex (M12 = 71 % of explained variance).

**Table 1:**
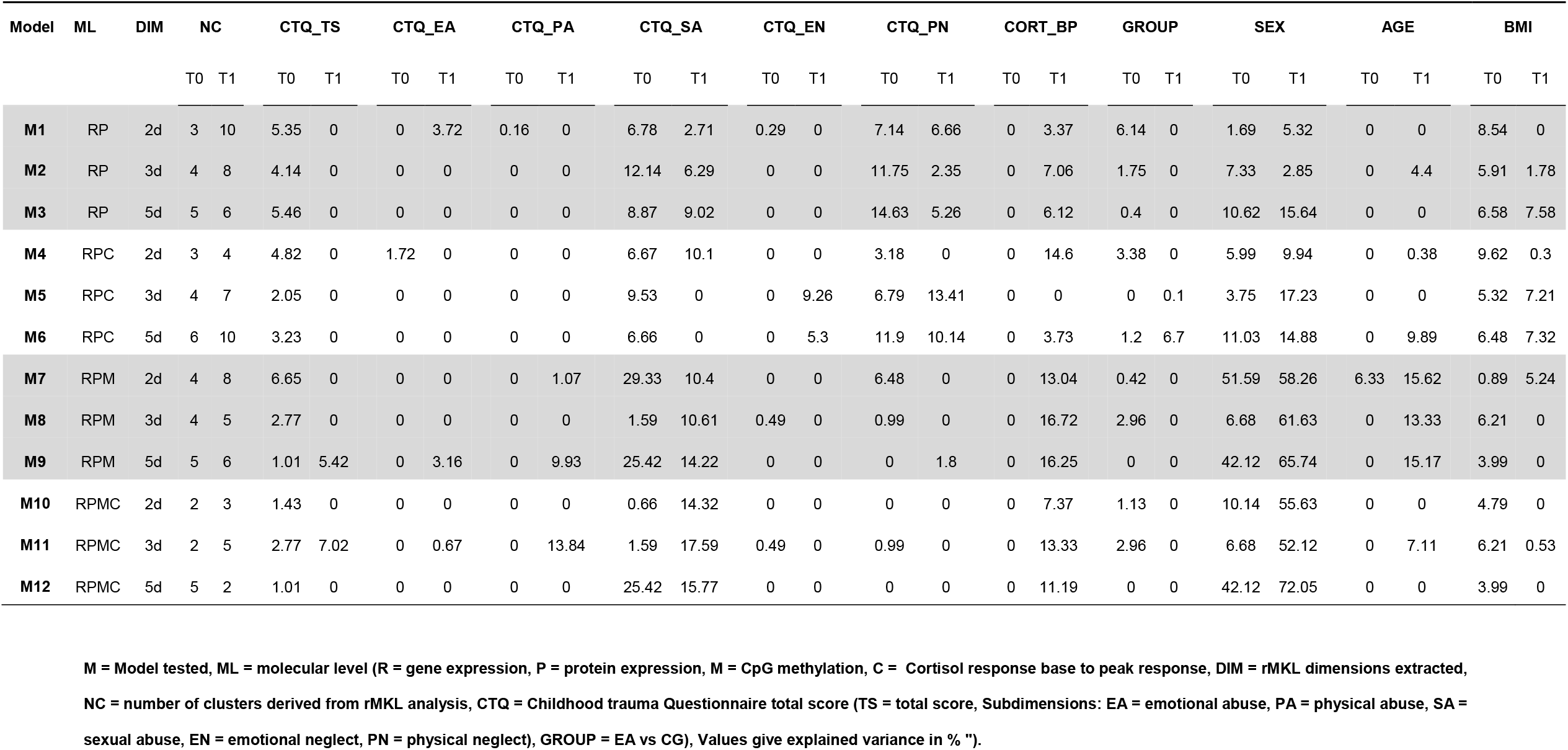
Variance explained by baseline and post-stress rMKL-models.

| Model | ML | DIM | NC |  | CTQ_TS |  | CTQ_EA |  | CTQ_PA |  | CTQ_SA |  | CTQ_EN |  | CTQ_PN |  | CORT_BP |  | GROUP |  | SEX |  | AGE |  | BMI |  |
| --- | --- | --- | --- | --- | --- | --- | --- | --- | --- | --- | --- | --- | --- | --- | --- | --- | --- | --- | --- | --- | --- | --- | --- | --- | --- | --- |
|  |  |  | T0 | T1 | T0 | T1 | T0 | T1 | T0 | T1 | T0 | T1 | T0 | T1 | T0 | T1 | T0 | T1 | T0 | T1 | T0 | T1 | T0 | T1 | T0 | T1 |
| M1 | RP | 2d | 3 | 10 | 5.35 | 0 | 0 | 3.72 | 0.16 | 0 | 6.78 | 2.71 | 0.29 | 0 | 7.14 | 6.66 | 0 | 3.37 | 6.14 | 0 | 1.69 | 5.32 | 0 | 0 | 8.54 | 0 |
| M2 | RP | 3d | 4 | 8 | 4.14 | 0 | 0 | 0 | 0 | 0 | 12.14 | 6.29 | 0 | 0 | 11.75 | 2.35 | 0 | 7.06 | 1.75 | 0 | 7.33 | 2.85 | 0 | 4.4 | 5.91 | 1.78 |
| M3 | RP | 5d | 5 | 6 | 5.46 | 0 | 0 | 0 | 0 | 0 | 8.87 | 9.02 | 0 | 0 | 14.63 | 5.26 | 0 | 6.12 | 0.4 | 0 | 10.62 | 15.64 | 0 | 0 | 6.58 | 7.58 |
| M4 | RPC | 2d | 3 | 4 | 4.82 | 0 | 1.72 | 0 | 0 | 0 | 6.67 | 10.1 | 0 | 0 | 3.18 | 0 | 0 | 14.6 | 3.38 | 0 | 5.99 | 9.94 | 0 | 0.38 | 9.62 | 0.3 |
| M5 | RPC | 3d | 4 | 7 | 2.05 | 0 | 0 | 0 | 0 | 0 | 9.53 | 0 | 0 | 9.26 | 6.79 | 13.41 | 0 | 0 | 0 | 0.1 | 3.75 | 17.23 | 0 | 0 | 5.32 | 7.21 |
| M6 | RPC | 5d | 6 | 10 | 3.23 | 0 | 0 | 0 | 0 | 0 | 6.66 | 0 | 0 | 5.3 | 11.9 | 10.14 | 0 | 3.73 | 1.2 | 6.7 | 11.03 | 14.88 | 0 | 9.89 | 6.48 | 7.32 |
| M7 | RPM | 2d | 4 | 8 | 6.65 | 0 | 0 | 0 | 0 | 1.07 | 29.33 | 10.4 | 0 | 0 | 6.48 | 0 | 0 | 13.04 | 0.42 | 0 | 51.59 | 58.26 | 6.33 | 15.62 | 0.89 | 5.24 |
| M8 | RPM | 3d | 4 | 5 | 2.77 | 0 | 0 | 0 | 0 | 0 | 1.59 | 10.61 | 0.49 | 0 | 0.99 | 0 | 0 | 16.72 | 2.96 | 0 | 6.68 | 61.63 | 0 | 13.33 | 6.21 | 0 |
| M9 | RPM | 5d | 5 | 6 | 1.01 | 5.42 | 0 | 3.16 | 0 | 9.93 | 25.42 | 14.22 | 0 | 0 | 0 | 1.8 | 0 | 16.25 | 0 | 0 | 42.12 | 65.74 | 0 | 15.17 | 3.99 | 0 |
| M10 | RPMC | 2d | 2 | 3 | 1.43 | 0 | 0 | 0 | 0 | 0 | 0.66 | 14.32 | 0 | 0 | 0 | 0 | 0 | 7.37 | 1.13 | 0 | 10.14 | 55.63 | 0 | 0 | 4.79 | 0 |
| M11 | RPMC | 3d | 2 | 5 | 2.77 | 7.02 | 0 | 0.67 | 0 | 13.84 | 1.59 | 17.59 | 0.49 | 0 | 0.99 | 0 | 0 | 13.33 | 2.96 | 0 | 6.68 | 52.12 | 0 | 7.11 | 6.21 | 0.53 |
| M12 | RPMC | 5d | 5 | 2 | 1.01 | 0 | 0 | 0 | 0 | 0 | 25.42 | 15.77 | 0 | 0 | 0 | 0 | 0 | 11.19 | 0 | 0 | 42.12 | 72.05 | 0 | 0 | 3.99 | 0 |

M = Model tested, ML = molecular level (R = gene expression, P = protein expression, M = CpG methylation, C = Cortisol response base to peak response, DIM = rMKL dimensions extracted, NC = number of clusters derived from rMKL analysis, CTQ = Childhood trauma Questionnaire total score (TS = total score, Subdimensions: EA = emotional abuse, PA = physical abuse, SA = sexual abuse, EN = emotional neglect, PN = physical neglect), GROUP = EA vs CG), Values give explained variance in % ").

### A supervised pathway to integration: The Data Integration Analysis for Biomarker discovery using Latent variable approaches for Omics studies (DIABLO) approach

We fit multiple sparse partial least squares regression (sPLS) models and DIABLO models (block sPLS) to single omics (R, M, P) or multi omics (RP, RMP) datasets to investigate the added value of multi-omics integration for the models performance in differentiating between conditions (EA/CG). Assessment of the model’s performance based on BER comparison yielded two central insights. First, gene expression (single level) sPLS models clearly outperformed all other models and differentiated better between groups than models based on protein expression, CpG methylation or models integrating multiple omics-datasets. Second, models based on post stress analytes differentiated better between groups than baseline models and this was true for all single or multi-omics models (Figure S9). In tendency, model performance did not increase with the number of analytes and models comprising 20 (RPT, RPM & RP_T0) or 30 (RP_T1) analytes of each molecular level reached lowest error rates.

The top performing model was the post-stress gene expression model (R_T1_20) based upon 20 transcripts (Figure 2A). 13 of these transcripts are shared with the best performing baseline gene expression model (Figure S10, Table S10), 14 were shared with in the best performing post-stress RP model (Figure 2B) and none were shared with analytes included in the best performing post-stress RPM model (Figure 2C). Two of the R_T1_20 transcripts corresponded to analytes included in proteomic dataset (HSPA8, H1FX) and 16 CpGs map to the same gene names.

**Figure 2:**
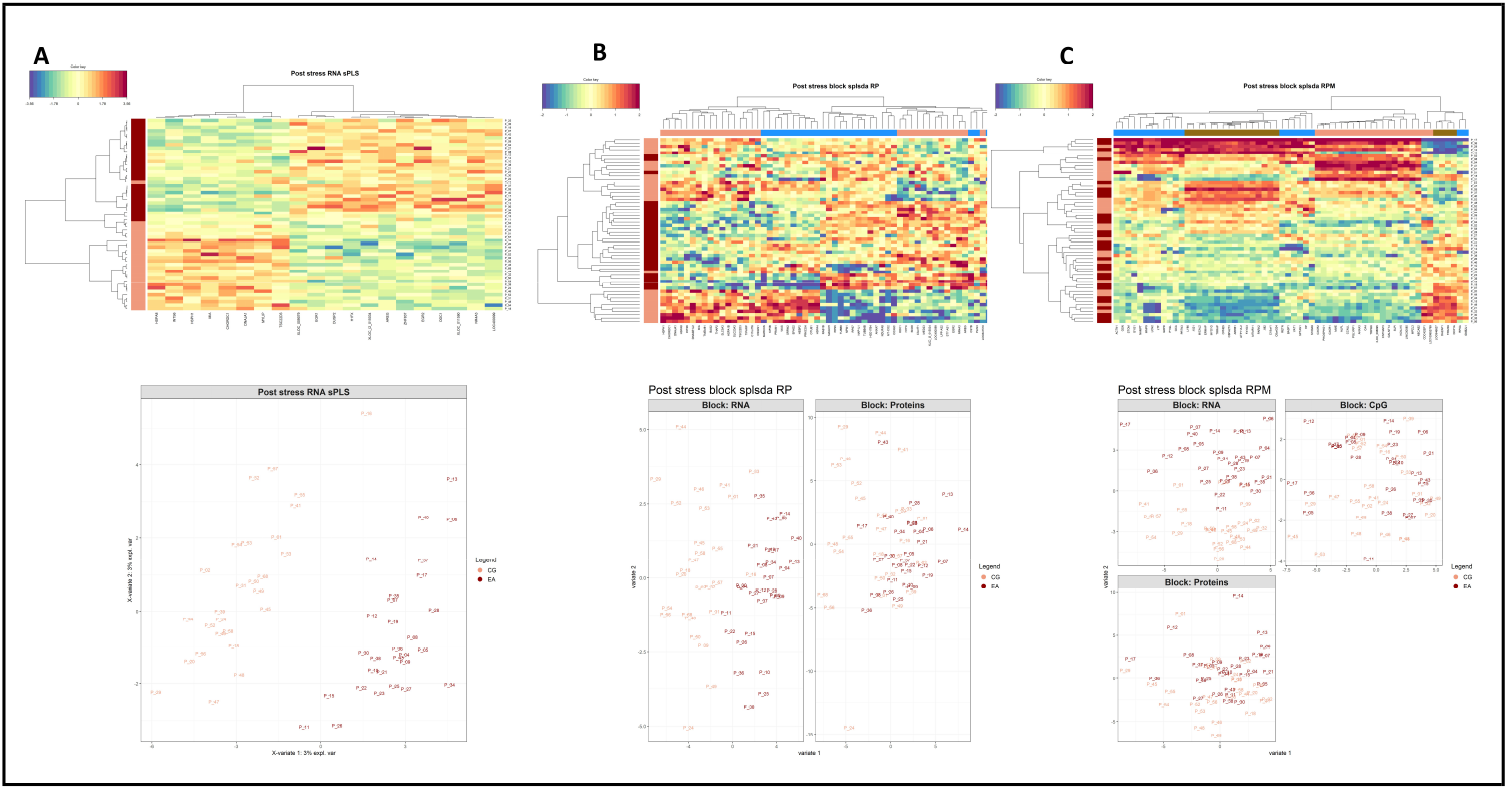
The model with the best discriminant capabilities (A) is based on 20 transcripts derived from CD14^+^ monocytes isolated 180 min after stress exposure. Including post stress proteomic data (B) and CpG methylation data did not result in a variable selection that differentiates better between participants’ with a history of childhood adversity and control participants. Heatmaps visualizes clustered analytes (columns) and participants (rows). Row annotations indicate participants’ condition (EA = dark red). Scatterplots visualize participants’ loadings (EA = dark red) on the first component extracted from analytes of included omic levels (A = RNA, B = RNA & Protein, C = RNA, Protein and CpG Methylation).

Cluster analyses clearly showed that the R_T1_20 model differentiates between participants based on two clusters of transcripts (Figure 2A). The first cluster contained transcripts downregulated (n = 8) and the second cluster contained transcripts upregulated (n = 12) in participants with a history of childhood adversity (Table 2). When supplied to the STRING database, downregulated transcripts formed a small network structure enriched for analytes functionally involved in Chaperone-mediated protein folding (GO:0061077), regulation of cellular response to heat (GO:1900034) and protein folding (GO:0006457) such as *HSPA8, HSPH1, CHORDC1 or DNAJA1*.

**Table 2:** Analytes of the best performing supervised model R_T1_20

| Characteristic |  | Overall, N = 56 | CG, N = 27 | EA, N = 29 | Difference<sup>1</sup> | 95% CI<sup>1,2</sup> | p-value<sup>1</sup> | q-value<sup>3</sup> |
| --- | --- | --- | --- | --- | --- | --- | --- | --- |
| HSPA8 | Mean (SD) | 13.26 (0.31) | 13.47 (0.24) | 13.06 (0.21) | 0.42 | 0.29, 0.54 | <0.001 | <0.001 |
| INTS6 | Mean (SD) | 8.28 (0.25) | 8.46 (0.16) | 8.11 (0.19) | 0.35 | 0.26, 0.44 | <0.001 | <0.001 |
| HSPH1 | Mean (SD) | 11.44 (0.38) | 11.72 (0.33) | 11.18 (0.20) | 0.53 | 0.38, 0.68 | <0.001 | <0.001 |
| MIA | Mean (SD) | 6.88 (0.47) | 7.19 (0.38) | 6.58 (0.33) | 0.61 | 0.42, 0.80 | <0.001 | <0.001 |
| CHORDC1 | Mean (SD) | 9.23 (0.26) | 9.43 (0.19) | 9.04 (0.15) | 0.39 | 0.30, 0.48 | <0.001 | <0.001 |
| DNAJA1 | Mean (SD) | 12.35 (0.28) | 12.57 (0.21) | 12.15 (0.14) | 0.42 | 0.32, 0.52 | <0.001 | <0.001 |
| MYLIP | Mean (SD) | 11.11 (0.32) | 11.32 (0.21) | 10.92 (0.29) | 0.40 | 0.26, 0.53 | <0.001 | <0.001 |
| TSC22D3 | Mean (SD) | 14.59 (0.35) | 14.82 (0.31) | 14.38 (0.24) | 0.43 | 0.29, 0.58 | <0.001 | <0.001 |
| XLOC_009079 | Mean (SD) | 8.79 (0.26) | 8.62 (0.20) | 8.95 (0.20) | -0.33 | -0.44, -0.22 | <0.001 | <0.001 |
| EGR1 | Mean (SD) | 10.13 (0.94) | 9.49 (0.64) | 10.72 (0.78) | -1.2 | -1.6, -0.85 | <0.001 | <0.001 |
| DUSP2 | Mean (SD) | 6.95 (0.53) | 6.59 (0.39) | 7.29 (0.42) | -0.70 | -0.92, -0.49 | <0.001 | <0.001 |
| H1FX | Mean (SD) | 11.03 (0.33) | 10.78 (0.27) | 11.26 (0.18) | -0.49 | -0.61, -0.36 | <0.001 | <0.001 |
| XLOC_l2_015034 | Mean (SD) | 6.43 (0.48) | 6.07 (0.36) | 6.78 (0.29) | -0.71 | -0.89, -0.53 | <0.001 | <0.001 |
| AREG | Mean (SD) | 4.63 (1.84) | 3.31 (0.79) | 5.85 (1.69) | -2.5 | -3.3, -1.8 | <0.001 | <0.001 |
| ZNF837 | Mean (SD) | 8.81 (0.37) | 8.55 (0.29) | 9.05 (0.26) | -0.50 | -0.65, -0.35 | <0.001 | <0.001 |
| EGR2 | Mean (SD) | 7.54 (1.00) | 6.84 (0.57) | 8.19 (0.87) | -1.3 | -1.7, -1.0 | <0.001 | <0.001 |
| ODC1 | Mean (SD) | 11.25 (0.39) | 10.97 (0.29) | 11.50 (0.29) | -0.53 | -0.68, -0.37 | <0.001 | <0.001 |
| XLOC_011590 | Mean (SD) | 5.82 (0.47) | 5.48 (0.37) | 6.13 (0.32) | -0.65 | -0.84, -0.46 | <0.001 | <0.001 |
| NR4A3 | Mean (SD) | 4.10 (1.76) | 2.77 (0.70) | 5.35 (1.51) | -2.6 | -3.2, -2.0 | <0.001 | <0.001 |
| LOC400099 | Mean (SD) | 10.67 (0.25) | 10.50 (0.20) | 10.83 (0.18) | -0.33 | -0.43, -0.23 | <0.001 | <0.001 |

<sup>1</sup> Welch Two Sample t-test
<sup>2</sup> CI = Confidence Interval
<sup>3</sup> False discovery rate correction for multiple testing

5 of the 12 transcripts upregulated in participants with a history of childhood adversity formed an interconnected network structure when supplied to the STRING Database. These transcripts include EGR1 and its paralog EGR2 as transcription factors crucially implicated in monocyte to macrophage differentiation and macrophage inflammatory functioning DUSP2, a member of the dual-specificity phosphatase family which is among others involved in immune cell activation, TSC22D3, a glucocorticoid-induced leucine zipper involved in monocyte anti-inflammatory signaling and NR4A3, a transcription factor causing the differentiation of monocytes to dendritic cells following exposure to inflammatory signaling

## Discussion

In the investigation of gene-environment interplay and the molecular signatures underlying stress related mediation of environmental risk, systems biology-oriented approaches have been proposed to gain a more comprehensive understanding of the complex multiple interrelations both within and between the different levels of analysis. Here, we utilised three approaches of integrating transcriptomic, proteomic, and CpG methylation data to test whether such integration can provide meaningful clues to biological alterations associated with the experience of early adversity, and whether combining multiple levels of analyses enhances group discrimination based on biological measures. Omics-data derive from cells isolated before and after stress exposure, based on the rationale that alterations in stress-associated molecular dynamics should be observable more clearly after stimulation. Following the three outlined paths to integration, we gained the following insights:

First, co-expression network analysis on single-omics level resulted in the identification of protein, gene co-expression and CpG co-methylation modules. Using the modules eigenvalues, a cross-omics correlational network was constructed, to test associations between modules and associations with variables of interest, such as history of early adversity or the cortisol stress response. Overall, correlation between identified modules was moderate, with occasional substantial correlations, especially between modules of the same analytical origin. The topographical representation of cross-omics interrelation reflected these observations, as co-expression modules of the same analytic origin clustered together. Furthermore, gene expression modules and early adversity clustered closely together, followed by protein modules, which mirrors the observation of significant associations between several gene and protein co-expression modules with childhood adversity. These modules were among others enriched for transcripts and proteins implicated in immune system related processes, or mitochondrial functioning. Interestingly, none of the co-methylation modules were associated with early adversity. Overall, there was a negative correlation between DNA methylation and gene expression modules.

Second, the joint analysis of different omics datasets with unsupervised and supervised integrative approaches yielded mixed results. In unsupervised analyses, especially those clusters that included three- or four-level data (RNA, protein and DNA methylation, and cortisol) discriminated participants on the basis of sex, and also explained a substantial proportion of BMI variance. Overall, the different cluster solutions were not strongly related to participants history of childhood adversity. One exception was the above mentioned cluster, which not only predicted participants sex but were also related to the “sexual abuse” CTQ subscale. Most likely, this can be explained by increased number of females reporting sexual abuse. These results suggest that biological meaningful subgrouping can be achieved (here: biological sex), and that models based on more input layers outperform those based on fewer layers, but it also shows that these analysis did not discriminate between participants with or without reported childhood adversity.

Third, the discriminatory performance of supervised approaches did not profit from multi-omics integration. Including protein-expression and CpG methylation to single layer gene-expression models resulted in decreased capacity to differentiate between participants with a history of childhood adversity and control participants. The model with the best discriminatory performance was based on 20 transcripts measured after stress exposure and this single layer post-stress model outperformed all other protein or CpG methylation-based single layers modules as well. These findings support the results from cross-omics co-expression network analysis and our previously reported results (18), which showed a strong relationship between gene co-expression and early adversity.

We conclude, that - at least in our specific case - integration of different layers of omics data does not necessarily lead to better results compared to in-depth single layer analysis or single-omic based discriminatory analyses. From cross-omics network analyses, we did learn that gene co-expression pattern, and to a lesser degree, protein co-expression pattern, were associated with early adversity. In line with our previous single layer analyses, adversity related co-expression modules were among others functionally implicated in stress-immune interplay processes and mitochondrial biology. Supervised integrative approaches showed that a set of 20 transcripts differentiated clearly between participants with early adversity and controls. Of note, theses transcripts contained HSP70, a key regulator of glucocorticoid receptor activity, and transcripts involved in monocyte inflammatory function and might be interesting candidates in search for potential biomarkers or childhood adversity associated gene expression signatures. However, single layer input outperformed multi-layer input, and we thus did not observe a clear analytical gain through multi-omics integration here.

Several reasons relating to theoretical considerations, study design as well as data input or preprocessing might account for these findings.

First, multi-omics approaches are conceptionally intriguing and there is some logic to the assumption that combining different omic levels analytically results in a deeper understanding of the molecular signatures underlying a distinct phenotype (26).

Yet, it is early days for these kind of analyses and currently more and more methods for integration are being developed. Clinical application and evaluation over time will give a clearer picture of what kind of biological and clinical data benefit from integration using specific methods or a combination of integration approaches. The fact that the discriminatory analyses we conducted here did not benefit from integration of multiple omic levels does not necessarily apply to the application of other approaches running on different statistics with a distinct analytical focus. Yet, in terms of parsimony, an important result derived from multi-omics approaches can be that that performing an integrational multi omics analysis might not be necessary in first place and such a finding could especially be of value in the context of biomarker development.

Second, although we assessed DNA methylation and protein abundance, our study design will most likely have favoured and thus prioritised mRNA expression as the most relevant marker. Given the well-established link between childhood adversity and altered stress system regulation, the rationale of the study was to investigate cellular adaptions following acute stress exposure. The observed adversity-related differences in gene expression might overshadow effects of other layers particularly in light of the number of different analytes (see below). More fine-grained time series in the context of stress exposure as well as time-lagged cross-omics correlations should be incorporated in future studies. On a more technical note, we used established pipelines to pre-process our data prior to analysis. While these steps are fitted to each layer specifically and prepare the data types for later on analysis, they do not correspond to each other so that some cross-omic signatures might have been lost during theses processes.

Third, there is an imperfect match in number and coverage between our analytes. Whereas mRNA transcripts cover a great majority of the transcriptome, the 450.000 investigated CpGs cover only a fraction of DNA methylation in the genome. On the level of protein, 1119 analytes included also only represent a fraction of the monocyte proteome. For instance, of the 20 transcripts differentiating between groups only two are represented within the protein dataset, so it is unclear whether these differences would amplify on the actual protein level. To make use the full range of available biological information we supplied all identified analytes to the integrational approaches applied instead of running analyses on the restricted but intersecting dataset of altogether 1065 analytes that mapped to the same gene symbol across omic levels. While this approach maximises information depth, individual omic levels are also weighted more heavily according to the number of analytes they contain, potentially overshadowing the information content of other omics levels. Future studies should aim at integrating “true” genome-wide and proteome-wide data.

To conclude, although we see promise in adopting multi-omics approaches in a systems biology framework to advance our understanding of molecular and gene regulatory processes targeted by early adversity our results were mixed here. We found some evidence for omic informed clustering, and unsupervised analyses differentiated participants according to biological variables, for example biological sex. Stress related gene-expression pattern were most strongly associated with childhood adversity, and integrating multiple cellular layers within an supervised approach did not result in better discriminatory performance. Our results emphasize the importance of carefully conducted single layer analyses and they as well underscore the importance of analysing molecular consequences of childhood adversity within the context of stress related physiological activation.

## References

1. Sonuga-Barke E, Kennedy M, Kumsta R, Knights N, Golm D, Rutter M, et al. Child-to-adult neurodevelopmental and mental health trajectories after early life deprivation: the young adult follow-up of the longitudinal English and Romanian Adoptees study.Lancet. 2017;389(10078):1539–48.

2. Teicher MH, Gordon JB, Nemeroff CB. Recognizing the importance of childhood maltreatment as a critical factor in psychiatric diagnoses, treatment, research, prevention, and education. Molecular psychiatry. 2022;27(3):1331–8.

3. Snyder-Mackler N, Burger JR, Gaydosh L, Belsky DW, Noppert GA, Campos FA, et al. Social determinants of health and survival in humans and other animals. Science. 2020;368(6493).

4. Lupien SJ, McEwen BS, Gunnar MR, Heim C. Effects of stress throughout the lifespan on the brain, behaviour and cognition. Nat Rev Neurosci. 2009;10(6):434–45.

5. Heim CM, Entringer S, Buss C. Translating basic research knowledge on the biological embedding of early-life stress into novel approaches for the developmental programming of lifelong health. Psychoneuroendocrinology. 2019;105:123–37.

6. Houtepen LC, Vinkers CH, Carrillo-Roa T, Hiemstra M, van Lier PA, Meeus W, et al. Genome-wide DNA methylation levels and altered cortisol stress reactivity following childhood trauma in humans. Nat Commun. 2016;7:10967.

7. Cole SW. Human social genomics. PLoS Genet. 2014;10(8):e1004601.

8. Haas R, Zelezniak A, Iacovacci J, Kamrad S, Townsend S, Ralser M. Designing and interpreting ‘multi-omic’ experiments that may change our understanding of biology. Curr Opin Syst Biol. 2017;6:37–45.

9. Pinu FR, Beale DJ, Paten AM, Kouremenos K, Swarup S, Schirra HJ, et al. Systems Biology and Multi-Omics Integration: Viewpoints from the Metabolomics Research Community. Metabolites. 2019;9(4).

10. Yan J, Risacher SL, Shen L, Saykin AJ. Network approaches to systems biology analysis of complex disease: integrative methods for multi-omics data. Brief Bioinform. 2018;19(6):1370–81.

11. Krassowski M, Das V, Sahu SK, Misra BB. State of the Field in Multi-Omics Research: From Computational Needs to Data Mining and Sharing. Front Genet. 2020;11:610798.

12. Breen MS, Tylee DS, Maihofer AX, Neylan TC, Mehta D, Binder EB, et al. PTSD Blood Transcriptome Mega-Analysis: Shared Inflammatory Pathways across Biological Sex and Modes of Trauma. Neuropsychopharmacology. 2018;43(3):469–81.

13. Jamil IN, Remali J, Azizan KA, Nor Muhammad NA, Arita M, Goh HH, et al. Systematic Multi-Omics Integration (MOI) Approach in Plant Systems Biology. Front Plant Sci. 2020;11:944.

14. Subramanian I, Verma S, Kumar S, Jere A, Anamika K. Multi-omics Data Integration, Interpretation, and Its Application. Bioinform Biol Insights. 2020;14:1177932219899051.

15. Speicher NK, Pfeifer N. Integrating different data types by regularized unsupervised multiple kernel learning with application to cancer subtype discovery. Bioinformatics. 2015;31(12):i268–75.

16. Singh A, Shannon CP, Gautier B, Rohart F, Vacher M, Tebbutt SJ, et al. DIABLO: an integrative approach for identifying key molecular drivers from multi-omics assays. Bioinformatics. 2019;35(17):3055–62.

17. Shannon CP, Blimkie TM, Ben-Othman R, Gladish N, Amenyogbe N, Drissler S, et al. Multi-Omic Data Integration Allows Baseline Immune Signatures to Predict Hepatitis B Vaccine Response in a Small Cohort. Front Immunol. 2020;11:578801.

18. Schwaiger M, Grinberg M, Moser D, Zang JC, Heinrichs M, Hengstler JG, et al. Altered Stress-Induced Regulation of Genes in Monocytes in Adults with a History of Childhood Adversity. Neuropsychopharmacology. 2016;41(10):2530–40.

19. Frach L, Tierling S, Schwaiger M, Moser D, Heinrichs M, Hengstler JG, et al. The mediating role of KITLG DNA methylation in the association between childhood adversity and cortisol stress reactivity does not replicate in monocytes. Psychoneuroendocrinology. 2020;116:104653.

20. Zang JCS, May C, Hellwig B, Moser D, Hengstler JG, Cole S, et al. Proteome analysis of monocytes implicates altered mitochondrial biology in adults reporting adverse childhood experiences. Transl Psychiatry. 2023;13(1):31.

21. Marwick B, Krishnamoorthy K. cvequality: Tests for the Equality of Coefficients of Variation from Multiple Groups. R software package version 0.1.3 ed2019.

22. Hoffman GE, Schadt EE. variancePartition: interpreting drivers of variation in complex gene expression studies. BMC Bioinformatics. 2016;17(1):483.

23. Langfelder P, Horvath S. WGCNA: an R package for weighted correlation network analysis. BMC Bioinformatics. 2008;9:559.

24. Epskamp S, Cramer AOJ, Waldorp LJ, Schmittmann VD, Borsboom D. qgraph : Network Visualizations of Relationships in Psychometric Data. J Stat Softw. 2012;48(4).

25. Rohart F, Gautier B, Singh A, Le Cao KA. mixOmics: An R package for ‘omics feature selection and multiple data integration. PLoS Comput Biol. 2017;13(11):e1005752.

26. Sathyanarayanan A, Mueller TT, Ali Moni M, Schueler K, members ETN, Baune BT, et al. Multi-omics data integration methods and their applications in psychiatric disorders. Eur Neuropsychopharmacol. 2023;69:26–46.

