## Supplements for "Biological embedding of childhood adversity - a multi-omics perspective on stress regulation"

#### Study design and context

This study is based on a project investigating long-term consequences of childhood adversity on stress reactivity, which included the collection of transcriptomic, proteomic, and genome-wide DNA methylation data. Initial work focused on individual omics data and provided insights into childhood adversity and stress related effects at the DNA methylation (Frach et al., 2019), gene expression (Dieckmann, Cole, & Kumsta, 2020; Schwaiger et al., 2016) and protein expression (Zang et al.) level. Here we take an integrative state-of-the-art multi-omics approach to further characterize molecular signatures associated with adverse childhood and acute stress experience. The subsequent analyses incorporate gene and protein expression data derived from monocytes collected during a baseline measurement minutes 45 min. before (t0) and 180 min. following exposure to psychosocial stress (t1). Genome-wide DNA methylation data derives from monocytes collected 45 min, after stress exposure. Cortisol measures were taken before and after exposure to the Trier Social Stress Test (TSST), at -45, -2, 1, 10, 20, 30, 45 and 90 min relative to stress exposure.

#### Sample characteristics

Following data curation and preprocessing, samples from 29 participants with a history of childhood adversity (EA, 65.5 % female ) and 27 control participants (CG, 66.7% female) were included in the analysis. Mean age was $M$ = 52.38, $SD$ = 5.01 and mean BMI $M$ = 25.21, $SD$ = 4.29. Groups did not differ in age, BMI, resilience (RS-25) current psychopathological symptoms, further bio-behavioral, or blood parameters (Table S1). Participants with a history of childhood adversity achieved significant higher CTQ subscale and total scores, reported a higher number of previous mental disorders and exhibited an significant decreased cortisol response (base to peak) after exposure to the TSST $\beta$ = - 52.4, p<.01, $F(1,54)$ = 10.97, $p$<.01, $R^2$ = .15. Monocyte derived multi-omics datasets comprised altogether 504.819 analytes including expression values of 23.259 transcripts, 1.119 proteins, and methylation degrees of 480.441 CpGs. 1.065 of these analytes mapped to the same gene name and occasionally subsequent analyses were performed with reduced data sets to meet analytical requirements.

Table S1: Sample characteristics

| | | Early adversity group (n=29) | Control group (n = 27) | *P* |
|---|---|---|---|---|
| Age | mean ± SD | 52.93 ± 5.24 | 51.78 ± 4.79 | 0.39 |
| Sex | number of females | 19 (65.51) | 18 (66.67) | 1 |
| CTQ total score | mean ± SD | 65.76 ± 14.51 | 35.34 ± 5.93 | < .001 |
| CTQ categories scores | mean ± SD | | | |
| Sexual abuse | | 10.97 ± 7.12 | 5.48 ± 1.31 | < .001 |
| Physical abuse | | 10.93 ± 5.38 | 5.63 ± 1.01 | < .001 |
| Emotional abuse | | 14.97 ± 4.16 | 7.67 ± 2.99 | < .001 |
| Emotional neglect | | 18.17 ±3 .49 | 9.96 ± 3.11 | < .001 |
| Physical neglect | | 10.27 ± 2.75 | 6.56 ± 1.78 | < .001 |
| History of mental disorder | n (%) | 17 (58) | 7 (25) | 0.03 |
| Depression history | n (%) | 12 (41) | 3 (11) | 0.02 |
| BSI global severity index | mean ± SD | 0.46 ± 0.41 | 0.34 ± 0.32 | 0.20 |
| RS-25 total | mean ± SD | 136.4 ± 26.82 | 138.9 ± 25.71 | 0.61 |
| Body mass index | mean ± SD | 25.45 ± 4.77 | 24.96 ± 3.69 | 0.54 |
| Baseline blood count | | | | |
| Leukocytes | mean ± SD | 6.82 ± 1.6 | 6.63 ± 1.72 | 0.68 |
| Lymphocytes | mean ± SD | 30.29 ± 7.6 | 32.10 ± 6.97 | 0.36 |
| Monocytes | mean ± SD | 6.08 ± 1.6 | 6.77 ± 1.87) | 0.14 |
| Neutrophils | mean ± SD | 60.75± 7.8 | 57.78 ± 7.98 | 0.16 |
| MLR | mean ± SD | 5.33 ± 2.08 | 5.01 ± 1.41 | 0.51 |
| Flow cytometric control | | | | |
| -45 monocyte isolation | mean ± SD purity | 93.92 ± 5.0 | 91.33 ± 6.64 | 0.13 |
| +45 monocyte isolation | mean ± SD purity | 94.00 ± 6.22 | 91.25 ± 9.3 | 0.22 |
| +180 monocyte isolation | mean ± SD purity | 93.02 ± 9.46 | 92..96 ± 7.71 | 0.98 |

Abbreviations: BSI, Brief Symptom Inventory; RS-25, Resilience Scale, 25 item version. MLR: Monocyte to Lymphocyte ratio

#### Monocyte isolation

CD14+ Monocytes were isolated from 10 ml EDTA blood samples via immunomagnetic cell separation (MACS; Miltenyi Biotec, Germany). Fluorescence-activated cell sorting indicated high quality of isolated CD 14+ monocyte cell population (92.63 %). We found no evidence for group differences in baseline blood cell counts or monocyte isolation quality before or after stress exposure (Table S1). For each participant the monocyte-to-lymphocyte ratio (LMR) was calculated based on a standardized complete blood count established during an initial medical examination.

#### DNA methylation profiling and data preprocessing

Genomic DNA was extracted from monocytes and treated with sodium bisulfite by means of the EZ96 DNA Methylation Kit (Zymo Research) and according to the manufacturers' standard protocol. Methylation levels were quantified with single-nucleotide resolution using a HiScan®SQ system in combination with an Ilumina Infinium HumanMethylation450 BeadChip. Methylation profiling was conducted at the university of Saarbrücken and following condition specific sample randomization (EA/CG) to control for time series effects prior to analysis. Raw signal intensities were extracted by means of the Ilumina Genome Studio software. As a matter of quality control, the following parameters were applied to remove participants and CpG sites from the subsequent analysis. First, only measures from participants with less than 1% (detection pvalue >0.01) of missing methylation values across all CpG sites were included in the analysis and samples were removed if they appeared as outliers with the comparison of the non-normalized and eventually normalized data. CpGs were removed from analysis based on missing methylation values (detection p-value > 0.001 in > = 1% of samples), bead count (< 3 in >= 5% of samples), cross reactive probes or CpGs within a 10 bp proximity to the primer were excluded from the analysis. Detection pvalue calculation and subsequent filtering steps were carried out using the R minfi (version 1.36 ) and wateRmelon package (version 1.34). Methylation values of 480.441 CpGs from 58 paticipants were retained for further analysis. Qunatro analysis (package version 1.24) showed an global normalization approach as appropriate and BMIQ normalization was applied to all valuies by means of the wateRmelon package. M-values were calculated as $M=\log_{2(}Beta/(1-Beta))$ and due to the more suitable statistical characteristics, all subsequent statistical analysis were performed on m-values. Batch effects were investigated by visual MDS plot inspection and we accounted for array and position related batch effect using the ComBat (sva package, version 3.38) .

**Gene expression profiling and data preprocessing**

RNA was isolated from monocytes (Macherey-Nagel, Germany) and RNA integrity analysis was performed on a Agilent 2100 bioanalyzer (Agilent Technologies) and indicated an overall excelent quality of the extracted ribonucleic acid (RIN: M = 9.7, SE = .03). Genome-wide gene expression profiling was realized using 100ng of RNA and Agilent Whole Human Genome Oligo Microarrays 8x60K V2. Analysis were carried out at the MACS Molecular Service Center (Miltenyi Biotech, Germany) following the manufacturer`s standard protocol could be performed on altogether 118 samples (EA: n = 30, CG: n = 29). The R qunatro package (version 3.1.1) was used to test for the eligibility of global or time point specific normalization and quantile-normalization was performed across all samples before log2-transformation of the data. Relative log expression (RLE) plots were used to asses successful normalization. A first filtering step was applied to remove transcripts with low expression values. Median expression was calculated for baseline, post stress and the joint expression data. Visual inspection of density plots suggested a cut-off of 2.5 and all transcripts (n = 3.579) that fell below this cutoff in 80% of participants within the early adversity and the control group before (t0) and after stress exposure (t1) were removed from the dataset. Next, all transcripts (n= 1.741) that mapped to multiple gene symbols were removed, which resulted in a final dataset containing expression values of 23.259 transcripts. Finally, we used hierarchical cluster analysis with Euclidian distance and RLE inspection on the final dataset to identify potential sample outliers. Two control groups' samples and two early adversity group samples showed slightly higher variation and were not excluded but flagged for observation in the subsequent analyses.

**Proteome profiling and data preprocessing**

Protein was extracted from monocytes using repeated exposure to ultrasonic waves and the application of physical force under precooled and iced conditions. 5 µg of extracted proteins were tryptic digested and 450 ng of tryptic peptides used for the subsequent proteome analysis by Liquid Chromatography tandem Mass spectrometry (LC-MS/MS ) analysis. Concentration of extracted proteins/peptides was determined in duplicates by amino acid analysis. Proteome analysis was performed on 118 samples (EA: n = 30, CG: n = 29). We additionally measured 22 external (EC) and 14 sample-derived (SC) standards to control for performance stability and technical variance. In order to avoid time-series effects, samples were run in clusters of four accounted for participants' condition (EA/CG), sex (male/female) and point of measurement (t0/t1) and control samples were run before and after each cluster. All peptides were separated on an Ultimate 3000 RSLCnano LC-system (Dionex) coupled to a nano-electrospray ion source (Thermo Scientific). For this, peptide mixtures were loaded on a 2 cm Acclaim PepMAP ® 100 microcapillary column (Thermo Scientific) packed with 5 µm C18 resin followed by 50 cm Acclaim PepMAP ® RSLC microcapillary column (Thermo Scientific) packed with 2 µm C18 resin by an auto sampler at a flow rate of 30 µL/ min at 60 °C . Peptide were eluted using a 120 min gradient of 5–40% B gradient. Mobile phase solvents consisted of 95% a (0.1% formic acid) and 5% B (84% acetonitrile, 0.1% formic acid) and B was subsequently increased over running time. MS/MS analysis was performed on a Q Exactive hybrid quadrupole-Orbitrap mass spectrometer. One full MS scan (350–1,400 m/z) was acquired in the Orbitrap each data selection cycle with 70.000 resolution setting, an automatic gain control (AGC) setting of 3 x e6 and a maximum ion accumulation time of 80 ms.MS2 analysis was performed with a top 10 setting with high-energy collision induced dissociation (HCD) with a collision energy of 27%, an AGC setting of 1 x e6, an isolation window of 2,2 m/z and a maximum ion accumulation time of 120 ms. Analysis of fragment ions was performed in an Orbitrap analyses with an resolution of 35.000 at 200 m/z, target of 1e6 and an accumulation time of 120 ms. Process performance was controlled in real-time and Preview was used to identify y the most common peptide modifications presented in the samples. MS Raw data was further processes in Max Quant and normalization across all 118 samples was performed by application of the MaxLFQ algorithm. Around 3500 protein groups were identified by LC-MS/MS analysis and only proteins identified by at least two unique peptides with valid values in at least eighty percent of one of the compared groups (EA/CG) both before and following stress exposure where accepted. The final dataset comprised expression values of 1119 proteins. Missing values were imputed through a protein specific minimum and LFGQ normalized data was log2 –transformed prior to subsequent analysis.

**Co-expression and co-methylation network analysis**

Co-expression and co-methylation network analyses were constructed to examine correlation patterns given in the respective omics datasets using weighted correlation network analysis by means of the WGCNA R package (version 1.70-3 ) (Langfelder & Horvath, 2008). Gene co-expression and CpG co-methylation networks were constructed upon the top 10.000 analytes with the highest variability according to median absolute deviation (*MAD*). Protein co-expression analysis was calculated in the full protein dataset. For gene and protein expression levels, WGCNA analysis was performed on baseline and post stress measurements of gene and protein expression levels and on a joint dataset for the assessment of stress-related analyte co-expression (see supplements for details). Module eigengenes, eigenproteins and eigenCpGs (MEs) – the first component of each module representing most of the variation of analytes with the module - were calculated for each module and modules with highly correlated MEs ($r$ < .75) were merged before further analysis. We used module preservation analyses (Langfelder, Luo, Oldham, & Horvath, 2011). to investigate the extent to which modules identified in protein and gene co-expression networks are conserved within the stress network. *Z*summary values as a permutation based preservation statistic (> 10 indicates strong preservation) was calculated with 500 permutations for each module. Calculated MEs were further used to a) identify modules related to the experience of early adversity or relevant other variables (modules of interest) and b) to investigate cross-omics interrelation of identified co-expression modules. For a) we calculated Pearson's correlation between MEs and clinical variables and further investigated the amount of variance explained in MEs through variables of interest. For b), two correlation-based approaches were applied. First, we used HCA to identify clusters of modules and variables based on correlation coefficients previously calculated between modules and clinical variables. Analyses were performed on baseline and post-stress modules separately (respectively gene expression and protein expression modules + CpG modules) and on modules identified in the joint data WGCNA. Results were visualized through a Heatmap. Second, we constructed two undirected correlation networks using the R qgraph package (version 1.9) (Epskamp, Cramer, Waldorp, Schmittmann, & Borsboom, 2012) to visualize cross-omics module interrelation and module – variable interrelations in baseline or post stress gene and protein co-expression modules and co-methylation modules. Edges below a weight of 0.3 were removed from both networks.

#### Biological meaning

Gene-ontology (GO) enrichment analysis were performed utilizing Enrichr (Kuleshov et al., 2016). Additionally we supplied analytes included in the best performing mixOmics models to the STRING database (Franceschini et al., 2012) to investigate interaction networks. Further characteristics are derived from the Gene Cards suit (Stelzer et al., 2016)

#### An unsupervised pathway to integration. The regularized multiple kernel learning for dimensionality reduction analysis approach (rMKL-DR)

rMKL-DR was performed using Locality Preserving Projections (LPP) for dimensionality reduction as suggested by Speicher and Pfeifer (2015). brief, rMKL-LPP represents an multivariate approach to multi-omics data integration (Subramanian, Verma, Kumar, Jere, & Anamika, 2020). This method applies implicit feature mapping to kernelize data, weights resulting kernel matrices based on informational content and while correcting for overfitting, uses LPP as an unsupervised method to preserve distance structures across matrices when projections into a lower subspace are computed. As such, rMKL-LP allows for the integration and dimensional reduction of the input data simultaneously and was reported to yield stable results even in smaller datasets (Speicher & Pfeifer, 2015). Kernel matrices representing gene expression, protein expression, genome-wide CpG methylation and participants cortisol response were generated by means of the web-rMKL offline preprocessing tool (Röder, Kersten, Herr, Speicher, & Pfeifer, 2019) with the Gaussian radial basis function (RBF) for kernel calculation. Subsequently, rMKL-LPP analyses were conducted within four different clusters. Each cluster involved different molecular levels and within each cluster, the number of projection dimension was varied and set to two, three or five (default setting). All analyses were performed separately for baseline and post stress data. The first cluster (M1 - M3) integrates protein expression and gene expression data. The second cluster adds participants' base-to-peak cortisol response to protein expression and gene expression data (M4 - M6). The third cluster includes gene expression, protein expression, and methylation data (M7 - M9). The fourth cluster (M10 - M12) integrates all data sets. Across all analyses, the number of neighbors was hold at a constant of nine (default setting) and following the considerations of Speicher and Pfeifer (2015), the number of dimensions of the projection subspace was kept rather small to extract a moderate number of subgroups. Next, we used multiple regression modelling (see above) to delineate sources of variation of rMKL-derived subgroups.

**A supervised pathway to integration: The Data Integration Analysis for Biomarker discovery using Latent variable approaches for Omics studies (DIABLO) approach.**

We utilized supervised discriminant analyses as implemented in the R mixOmics package (version 6.16.3) (Rohart, Gautier, Singh, & Lê Cao, 2017) to identify analytes explaining participants group membership (EA/CG). Analyses were performed either on each of the omics datasets separately (R = Gene expression data, P = protein expression data, M = Methylation data) using sparse partial least squares regression discriminant analysis (sPLSDA), on gene and protein expression data (RP) or gene expression, protein expression and CpG methylation data (RPM) simultaneously using block sparse partial least squares regression discriminant analysis (block sPLSDA) as provided by the DIABLO framework (Singh et al., 2019). Both types of analyses are based on Partial Least Squares (PLS) regression, to model and increase variance and co-variance structures within and across provided datasets to select correlated and contrasting analytes, which can discriminate between groups of interest. (Singh et al. 2017 , Singh et al., 2019). Within the DIABLO approach, PLS discriminant analysis is used to maximize correlation between clusters of highly correlated analytes derived from an initial Sparse Generalized Canonical Correlation Analysis (sGCCA). All analyses including gene and protein expression data were performed separately on baseline and post stress data (T0/T1) to identify analytes associated with the experience of childhood adversity and investigate the influence of stress on the compatibility of the defined omics models to differentiate between conditions. To select the model with the best discriminant performance on basis of a) the molecular levels and b) the number of analytes included in the models we ran all constructed omics models (M, R_T0, R_T1, P_T0, P_T1, RP_T0, RP_T1, RPM_t0 & RPM_T1) ten times while increasing the number of selected analytes by 10 in each step up to a total of 100. Model performance was evaluated on basis of balanced error rate (BER, smaller = better) and we compared BER across all models to select the model that discriminated best between participants with a history of childhood adversity and control participants.

*Table S2:* Intercorrelation of analyte specific variation in EA individuals (n = 29).

| CV (1) | CV (2) | r | 95% CI | t(27) | p |
|---|---|---|---|---|---|
| P_T0 | P_T1 | 0.22 | [-0.16, 0.54] | 1.18 | 0.249 |
| P_T0 | R_T0 | -0.36 | [-0.64, 0.01] | -1.98 | 0.058 |
| P_T0 | R_T1 | -0.12 | [-0.47, 0.25] | -0.65 | 0.521 |
| P_T0 | M | 0.19 | [-0.19, 0.52] | 1.03 | 0.313 |
| P_T1 | R_T0 | 0.07 | [-0.30, 0.43] | 0.37 | 0.715 |
| P_T1 | R_T1 | -0.20 | [-0.53, 0.17] | -1.09 | 0.287 |
| P_T1 | M | 0.03 | [-0.34, 0.39] | 0.14 | 0.890 |
| R_T0 | R_T1 | -0.32 | [-0.61, 0.06] | -1.74 | 0.094 |
| R_T0 | M | -0.31 | [-0.61, 0.06] | -1.69 | 0.103 |

CV = Coefficient of variation, P = protein expression, R = gene expression level , M = degree of methylation, T0 = baseline measure 45 min. before stress exposure, T1 = post stress measure 180 min following stress exposure.

Table S3: Intercorrelation of analyte specific variation in control individuals (n = 27).

| CV (1) | CV (2) | r | 95% CI | t(25) | p |
|---|---|---|---|---|---|
| P_T0 | P_T1 | 0.73 | [ 0.48, 0.87] | 5.28 | < .001*** |
| P_T0 | R_T0 | -0.14 | [-0.49, 0.26] | -0.70 | 0.492 |
| P_T0 | R_T1 | -0.27 | [-0.59, 0.12] | -1.42 | 0.168 |
| P_T0 | M | -0.04 | [-0.41, 0.35] | -0.19 | 0.848 |
| P_T1 | R_T0 | -0.39 | [-0.67, -0.01] | -2.12 | 0.044* |
| P_T1 | R_T1 | 0.12 | [-0.27, 0.48] | 0.63 | 0.536 |
| P_T1 | M | 0.02 | [-0.36, 0.40] | 0.10 | 0.924 |
| R_T0 | R_T1 | -0.03 | [-0.40, 0.36] | -0.13 | 0.900 |

CV = Coefficient of variation, P = protein expression, R = gene expression , M = degree of methylation, T0 = baseline measure 45 min. before stress exposure, T1 = post stress measure 180 min following stress exposure, p =pvalue.

*Table S4:* Comparison of analyte specific CV differences between groups.

| CV | comp | D_AD | p |
|---|---|---|---|
| M | EA/CG | 0.0002087109 | .9884735 |
| R_T0 | EA/CG | 3.760986e-07 | .9995107 |
| R_T1 | EA/CG | .936755e-07 | .9993852 |
| P_T0 | EA/CG | 0.000523163 | .9817518 |
| P_T1 | EA/CG | 0.03248318 | .8569713 |

CV = Coefficient of variation, P = protein expression, R = gene expression , M = degree of methylation, T0 = baseline measure 45 min. before stress exposure, T1 = post stress measure 180 min following stress exposure, comp = comparisons, EA= early adversity group, CG = control group, D_AD = DAD (asymptotic x2 ) test statistic, p =pvalue

*Table S5:* Variance fractions in protein expression data explained by variables.

| Variable | CTQ model | | | | | | | CTQ subscale model | | | | | | |
|---|---|---|---|---|---|---|---|---|---|---|---|---|---|---|
| | Baseline (t0) | | | post stress (t1) | | | | Baseline (t0) | | | post stress (t1) | | | |
| | *M* | *SD* | *R* | *M* | *SD* | *R* | *d* M (t1-t0) | *M* | *SD* | *R* | *M* | *SD* | *R* | *d* M (t1-t0) |
| EA | 0,21 % | 1,66 % | 19,55 % | 0,23 % | 1,80 % | 25,64 % | 0,02 % | 0,10 % | 1,22 % | 18,69 % | 0,00 % | 0,00 % | 0,00 % | -0,10 % |
| SEX | 1,18 % | 3,04 % | 28,03 % | 0,55 % | 2,11 % | 35,99 % | -0,62 % | 2,33 % | 5,31 % | 37,58 % | 1,33 % | 3,88 % | 27,90 % | -1,00 % |
| BMI | 2,34 % | 2,98 % | 20,50 % | 2,37 % | 3,08 % | 31,33 % | 0,04 % | 1,65 % | 2,33 % | 23,55 % | 1,80 % | 2,45 % | 27,25 % | 0,15 % |
| AGE | 1,93 % | 2,53 % | 22,65 % | 1,63 % | 2,22 % | 21,63 % | -0,30 % | 1,40 % | 2,07 % | 18,03 % | 1,28 % | 1,86 % | 18,52 % | -0,11 % |
| LMR | 4,30 % | 5,19 % | 30,70 % | 2,36 % | 2,89 % | 20,17 % | -1,94 % | 3,58 % | 4,79 % | 31,67 % | 1,91 % | 2,49 % | 18,28 % | -1,67 % |
| RS25 | 3,18 % | 3,26 % | 20,72 % | 2,51 % | 2,81 % | 20,86 % | -0,67 % | 2,85 % | 2,99 % | 18,49 % | 1,75 % | 2,18 % | 17,35 % | -1,10 % |
| CTQ | 2,07 % | 2,65 % | 17,32 % | 2,77 % | 3,43 % | 22,92 % | 0,70 % | | | | | | | |
| CTQ_sa | | | | | | | | 1,96 % | 2,51 % | 24,97 % | 1,64 % | 2,30 % | 18,71 % | -0,33 % |
| CTQ_pa | | | | | | | | 2,01 % | 2,34 % | 21,17 % | 1,84 % | 2,17 % | 12,88 % | -0,17 % |
| CTQ_en | | | | | | | | 2,81 % | 3,16 % | 18,18 % | 2,66 % | 3,16 % | 21,54 % | -0,15 % |
| CTQ_pn | | | | | | | | 2,55 % | 3,16 % | 23,34 % | 3,85 % | 4,85 % | 27,22 % | 1,30 % |
| CTQ_ea | | | | | | | | 1,95 % | 2,18 % | 13,93 % | 2,44 % | 2,78 % | 16,02 % | 0,49 % |
| Residuals | 84,80 % | 9,33 % | 67,39 % | 87,57 % | 6,67 % | 49,23 % | 2,78 % | 76,82 % | 11,54 % | 69,39 % | 79,50 % | 8,79 % | 61,80 % | 2,68 % |

EA: early adversity, CTQ: Childhood Trauma Questionnaire total score (CTQ categories: sa = sexual abuse, pa = physical abuse, ea= emotional abuse, en = emotional neglect, pn = physical abuse), RS25: Resilience Scale 25 total score, BMI: body mass index, SEX: biological sex, AGE: age in years at participation, LMR: lymphocyte to monocyte ratio, R = maximum of range.

*Table S6:* Variance in gene expression data explained by variables in different models.

| Variable | CTQ model | | | | | | | CTQ subscale model | | | | | | |
|---|---|---|---|---|---|---|---|---|---|---|---|---|---|---|
| | Baseline (t0) | | | post stress (t1) | | | | Baseline (t0) | | | post stress (t1) | | | |
| | *M* | *SD* | *R* | *M* | *SD* | *R* | *d M* (t1-t0) | *M* | *SD* | *R* | *M* | *SD* | *R* | *d* M (t1-t0) |
| EA | 1,02 % | 4,97 % | 59,23 % | 0,89 % | 4,53 % | 66,25 % | -0,13 % | 1,23 % | 5,78 % | 61,80 % | 0,94 % | 4,90 % | 65,55 % | -0,29 % |
| SEX | 0,57 % | 2,07 % | 33,27 % | 0,62 % | 2,34 % | 37,34 % | 0,06 % | 0,49 % | 2,43 % | 39,20 % | 0,60 % | 2,52 % | 33,49 % | 0,11 % |
| BMI | 2,04 % | 2,67 % | 24,93 % | 1,64 % | 2,25 % | 23,04 % | -0,40 % | 1,50 % | 2,13 % | 21,06 % | 1,20 % | 1,79 % | 19,72 % | -0,30 % |
| AGE | 1,93 % | 2,59 % | 27,11 % | 2,03 % | 2,66 % | 26,51 % | 0,10 % | 1,51 % | 2,15 % | 24,21 % | 1,58 % | 2,29 % | 29,24 % | 0,07 % |
| LMR | 1,55 % | 2,16 % | 23,94 % | 1,77 % | 2,52 % | 39,87 % | 0,22 % | 1,16 % | 1,72 % | 21,79 % | 1,34 % | 2,03 % | 29,75 % | 0,18 % |
| RS25 | 2,01 % | 2,65 % | 30,59 % | 1,56 % | 2,15 % | 24,29 % | -0,46 % | 1,62 % | 2,27 % | 29,92 % | 1,20 % | 1,80 % | 23,38 % | -0,42 % |
| CTQ | 2,36 % | 3,39 % | 39,26 % | 2,76 % | 3,82 % | 40,57 % | 0,40 % | | | | | | | |
| CTQ_sa | | | | | | | | 1,82 % | 2,50 % | 29,34 % | 1,78 % | 2,49 % | 33,14 % | -0,04 % |
| CTQ_pa | | | | | | | | 2,08 % | 2,70 % | 34,84 % | 2,10 % | 2,71 % | 34,38 % | 0,02 % |
| CTQ_en | | | | | | | | 2,80 % | 3,37 % | 31,51 % | 2,96 % | 3,57 % | 37,77 % | 0,16 % |
| CTQ_pn | | | | | | | | 2,03 % | 2,69 % | 26,96 % | 2,08 % | 2,88 % | 37,84 % | 0,05 % |
| CTQ_ea | | | | | | | | 2,44 % | 2,91 % | 25,79 % | 2,78 % | 3,24 % | 34,01 % | 0,34 % |
| Residuals | 88,52 % | 8,09 % | 71,31 % | 88,73 % | 8,06 % | 77,98 % | | 81,33 % | 9,62 % | 83,12 % | 81,45 % | 9,42 % | 84,20 % | |

EA: early adversity, CTQ: Childhood Trauma Questionnaire total score (CTQ categories: sa = sexual abuse, pa = physical abuse, ea= emotional abuse, en = emotional neglect, pn = physical abuse), RS25: Resilience Scale 25 total score, BMI: body mass index, SEX: biological sex, AGE: age in years at participation, LMR: lymphocyte to monocyte ratio, R = maximum of range.

*Table S7:* Variance in CpG methylation data explained by variables

| Variable | CTQ model | | | CTQ subscale model | | |
|---|---|---|---|---|---|---|
| | *M* | *SD* | *R* | *M* | *SD* | *R* |
| EA | 0,09 % | 1,18 % | 34,12 % | 0,11 % | 1,44 % | 37,54 % |
| SEX | 1,62 % | 8,58 % | 87,62 % | 1,59 % | 8,45 % | 87,41 % |
| BMI | 2,27 % | 3,09 % | 38,54 % | 1,50 % | 2,33 % | 36,62 % |
| AGE | 1,50 % | 2,03 % | 34,97 % | 1,05 % | 1,56 % | 27,49 % |
| LMR | 1,58 % | 2,13 % | 29,11 % | 1,17 % | 1,68 % | 26,36 % |
| RS25 | 1,54 % | 2,12 % | 28,73 % | 1,08 % | 1,63 % | 28,03 % |
| CTQ | 1,21 % | 1,90 % | 37,87 % | | | |
| CTQ_sa | | | | 1,66 % | 2,33 % | 34,90 % |
| CTQ_pa | | | | 1,88 % | 2,44 % | 28,76 % |
| CTQ_en | | | | 2,42 % | 2,86 % | 40,03 % |
| CTQ_pn | | | | 1,57 % | 2,09 % | 30,09 % |
| CTQ_ea | | | | 3,05 % | 3,49 % | 38,43 % |
| Residuals | 90,19 % | 9,68 % | 89,50 % | 82,90 % | 10,28 % | 89,30 % |

EA: early adversity, CTQ: Childhood Trauma Questionnaire total score (CTQ categories: sa = sexual abuse, pa = physical abuse, ea= emotional abuse, en = emotional neglect, pn = physical abuse), RS25: Resilience Scale 25 total score, BMI: body mass index, SEX: biological sex, AGE: age in years at participation, LMR: lymphocyte to monocyte ratio, R = maximum of range.

*Table S8:* Variance fractions in joint protein expression data explained by variables

| Variable | CTQ model | | | CTQ subscale model | | |
|---|---|---|---|---|---|---|
| | *M* | *SD* | *R* | *M* | *SD* | *R* |
| EA | 0,56 % | 2,34 % | 19,06 % | 0,20 % | 1,52 % | 19,38 % |
| EA:STRESS | 0,04 % | 0,32 % | 5,83 % | 0,04 % | 0,35 % | 6,28 % |
| SEX | 1,00 % | 2,39 % | 22,99 % | 2,05 % | 4,24 % | 25,51 % |
| STRESS | 0,15 % | 0,60 % | 7,71 % | 0,17 % | 0,63 % | 7,74 % |
| BMI | 1,82 % | 2,35 % | 20,93 % | 1,33 % | 1,78 % | 15,45 % |
| AGE | 1,36 % | 1,86 % | 16,77 % | 1,01 % | 1,54 % | 16,27 % |
| LMR | 2,52 % | 2,99 % | 18,74 % | 2,13 % | 2,79 % | 19,54 % |
| RS25 | 2,34 % | 2,40 % | 16,01 % | 2,00 % | 2,13 % | 14,51 % |
| CTQ | 2,03 % | 2,51 % | 15,06 % | | | |
| CTQ_sa | | | | 1,64 % | 2,10 % | 15,59 % |
| CTQ_pa | | | | 1,49 % | 1,63 % | 11,70 % |
| CTQ_en | | | | 2,12 % | 2,51 % | 18,35 % |
| CTQ_pn | | | | 2,50 % | 3,14 % | 19,97 % |
| CTQ_ea | | | | 1,37 % | 1,66 % | 10,81 % |
| Residuals | 88,18 % | 7,17 % | 54,90 % | 81,94 % | 9,09 % | 56,44 % |

EA: early adversity, CTQ: Childhood Trauma Questionnaire total score (CTQ categories: sa = sexual abuse, pa = physical abuse, ea= emotional abuse, en = emotional neglect, pn = physical abuse), RS25: Resilience Scale 25 total score, BMI: body mass index, SEX: biological sex, AGE: age in years at participation, LMR: lymphocyte to monocyte ratio, STRESS: baseline or post stress measurement, EA:STRESS = group (EA/CG) x STRESS interaction effect, R = maximum of range.

*Table S9:* Variance fractions in joint gene expression data explained by variables

| Variable | CTQ model | | | CTQ subscale model | | |
|---|---|---|---|---|---|---|
| | *M* | *SD* | *R* | *M* | *SD* | *R* |
| EA | 1,06 % | 4,44 % | 61,73 % | 1,20 % | 4,97 % | 63,55 % |
| EA:STRESS | 0,22 % | 0,98 % | 14,36 % | 0,22 % | 0,99 % | 16,03 % |
| SEX | 0,36 % | 1,54 % | 29,92 % | 0,36 % | 1,74 % | 32,03 % |
| STRESS | 0,39 % | 1,19 % | 18,34 % | 0,40 % | 1,20 % | 18,41 % |
| BMI | 0,90 % | 1,23 % | 11,48 % | 0,66 % | 0,98 % | 9,91 % |
| AGE | 0,89 % | 1,20 % | 13,48 % | 0,71 % | 1,02 % | 14,06 % |
| LMR | 0,86 % | 1,21 % | 15,42 % | 0,65 % | 0,98 % | 329,28 % |
| RS25 | 0,94 % | 1,24 % | 14,77 % | 0,74 % | 1,05 % | 273,67 % |
| CTQ | 1,50 % | 2,19 % | 25,19 % | | | |
| CTQ_sa | | | | 0,89 % | 1,26 % | 14,59 % |
| CTQ_pa | | | | 1,08 % | 1,38 % | 13,02 % |
| CTQ_en | | | | 1,58 % | 2,01 % | 23,19 % |
| CTQ_pn | | | | 1,11 % | 1,57 % | 22,10 % |
| CTQ_ea | | | | 1,38 % | 1,65 % | 13,57 % |
| Residuals | 92,87 % | 6,04 % | 67,11 % | 89,03 % | 7,12 % | 76,74 % |

EA: early adversity, CTQ: Childhood Trauma Questionnaire total score (CTQ categories: sa = sexual abuse, pa = physical abuse, ea= emotional abuse, en = emotional neglect, pn = physical abuse), RS25: Resilience Scale 25 total score, BMI: body mass index, SEX: biological sex, AGE: age in years at participation, LMR: lymphocyte to monocyte ratio), STRESS: baseline or post stress measurement, EA:STRESS = group (EA/CG) x STRESS interaction effect, R = maximum of range.

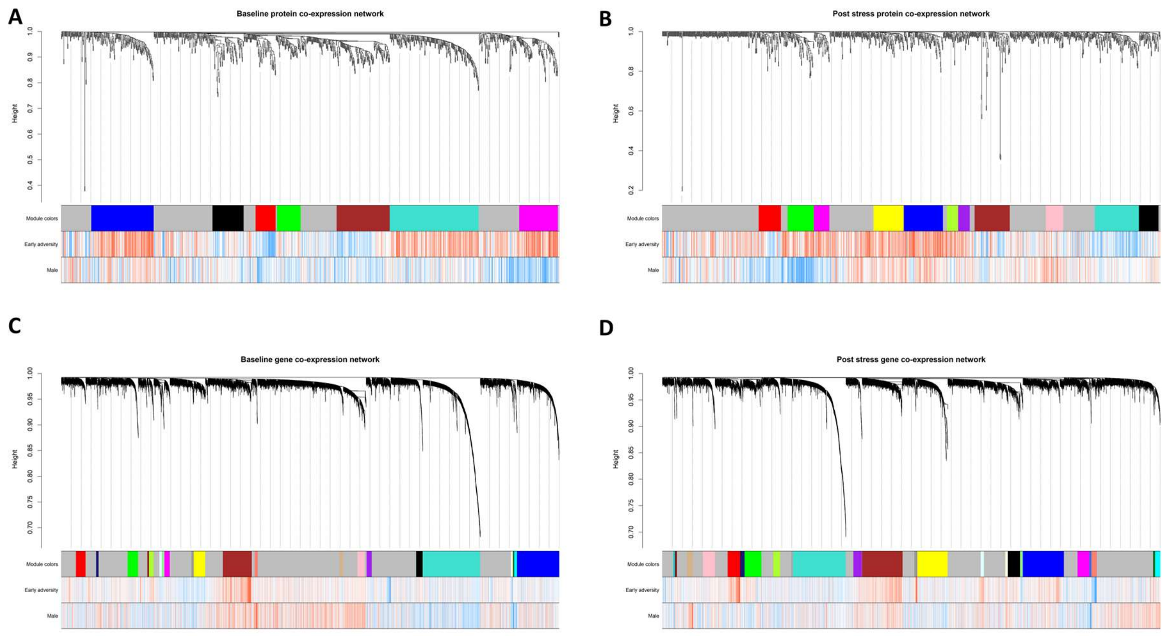


Figure S1. Dendrograms of the baseline (A) and post stress proteome (B) and the baseline (C) and post stress (D) transcriptome. Each line represents an analyte (leaf) and each low-hanging cluster represents a group of analytes with similar network connections (branch) on the tree. The first band underneath the tree indicates detected modules and subsequent bands indicate analyte-trait correlation. Red indicates a strong relationship and blue indicates a strong negative relationship.

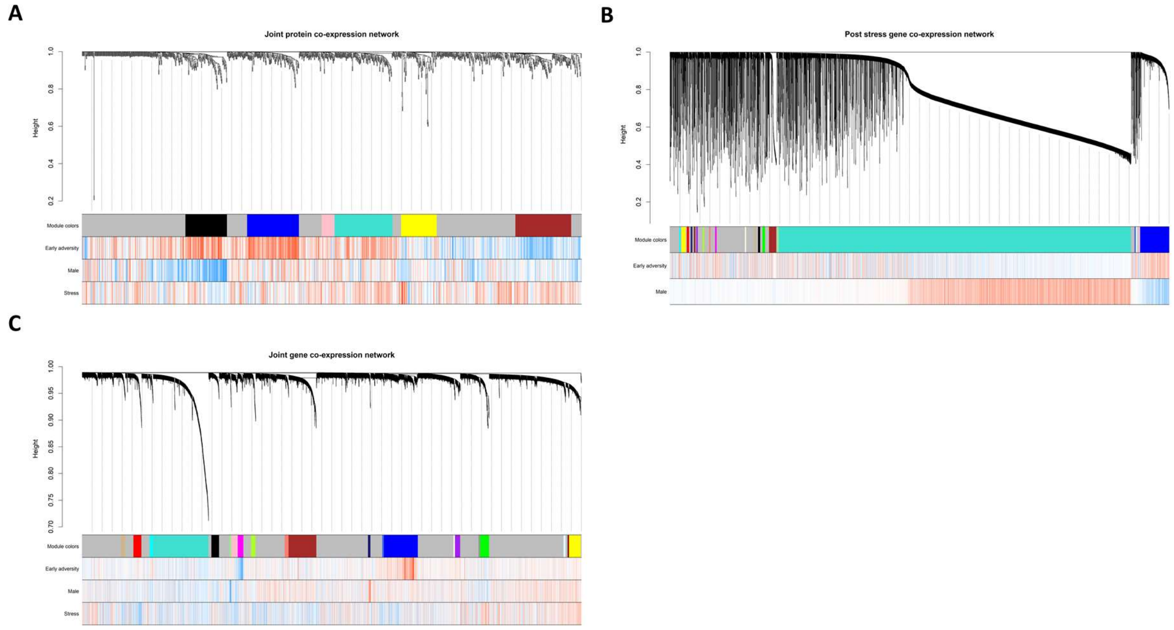


Figure S2. Dendrograms of the joint protein (A) and gene expression datasets (C) and the CpG methylation dataset (B). Each line represents an analyte (leaf) and each low-hanging cluster represents a group of analytes with similar network connections (branch) on the tree. The first band underneath the tree indicates detected modules and subsequent bands indicate analyte-trait correlation. Red indicates a strong relationship and blue indicates a strong negative relationship.

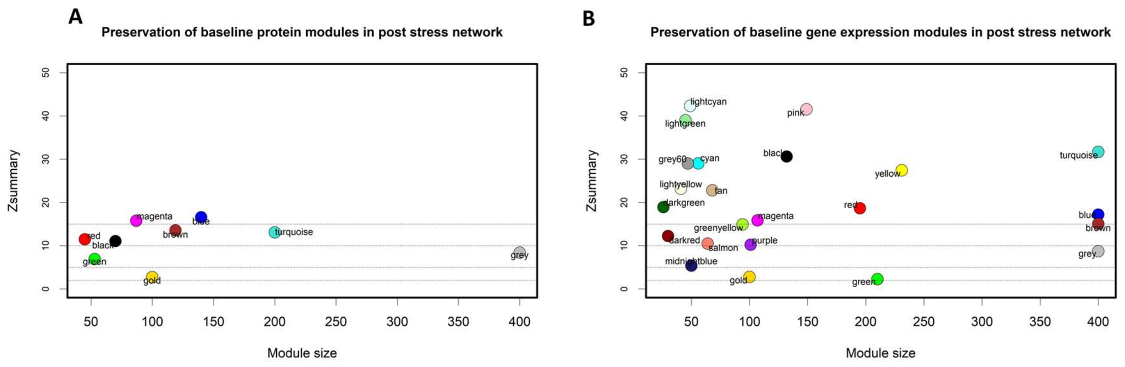


Figure S3. Preservation of baseline protein expression (A) and gene expression modules (B) in the post stress network is shown by plotting modules specific Zsummary scores against the module size. A Zsummary score < 2 indicates no module preservation, a score of 10 indicates weak preservation and a score > 10 indicates strong preservation of modules across psychotherapeutic intervention. Dashed lines indicate Zsummary values of 2, 5, 10 and 15. The gold module represents a random sample of the whole network; the grey module represents uncharacterized analytes not assigned to a module and as expected both, show low Zsummary scores.

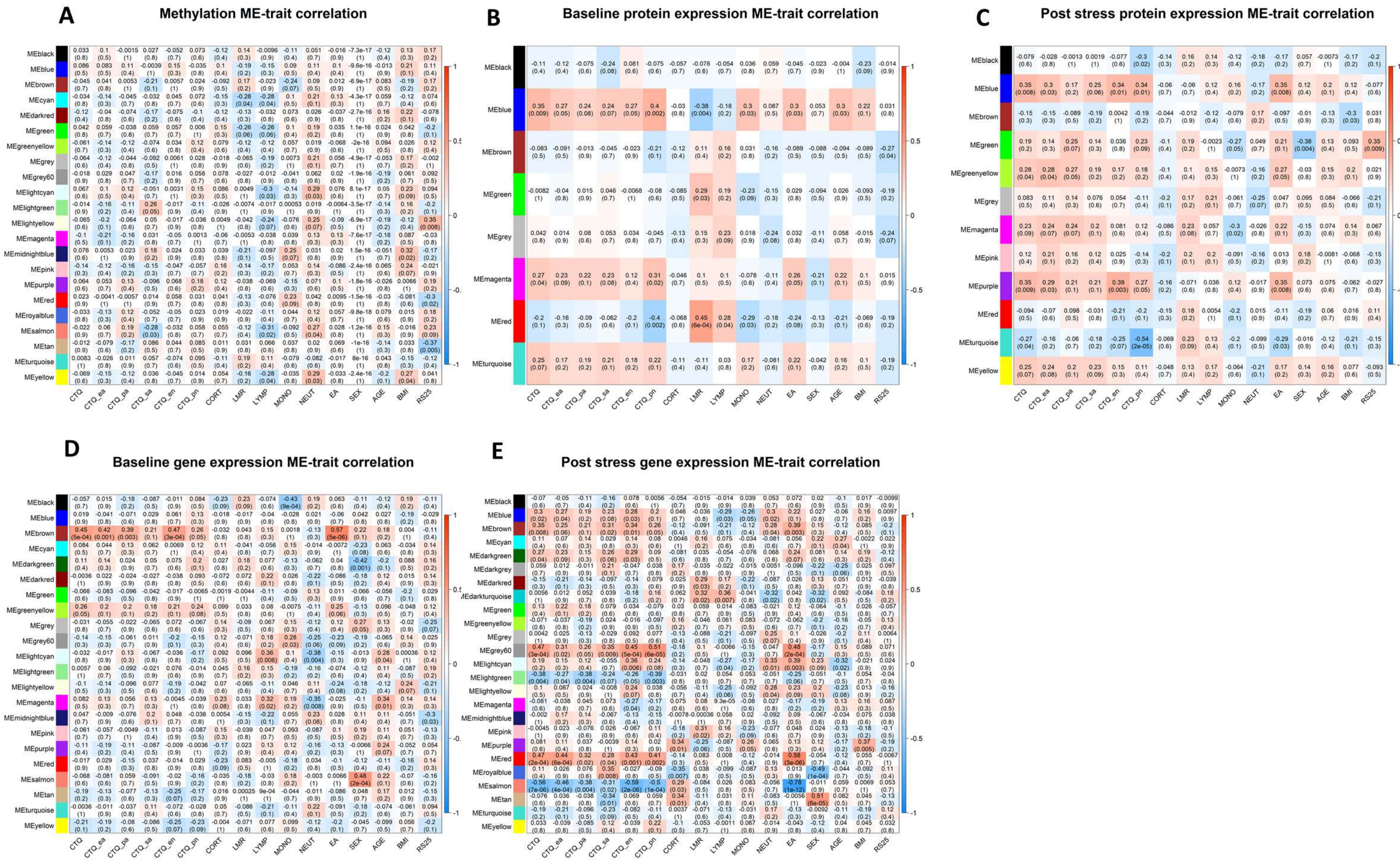


Figure S4. Module eigenproteins, eigengenes or eigenCpGs (ME) correlation with clinical variables are depicted for the CpG methylation network (A), the baseline and post-stress protein consensus network (B & C), the baseline and post-stress gene expression network (D & E). Abbreviations: EA: early adversity, CTQ: Childhood Trauma Questionnaire total score (CTQ categories: sa = sexual abuse, pa = physical abuse, ea= emotional abuse, en = emotional neglect, pn = physical abuse), RS25: Resilience Scale 25 total score, CORT: Cortisol base to peak ratio, BMI: body mass index, SEX: biological sex, AGE: age in years at participation, SMO: smoking, LYMPH & MONO: count of lymphocytes and monocytes, LMR: lymphocyte to monocyte ratio.

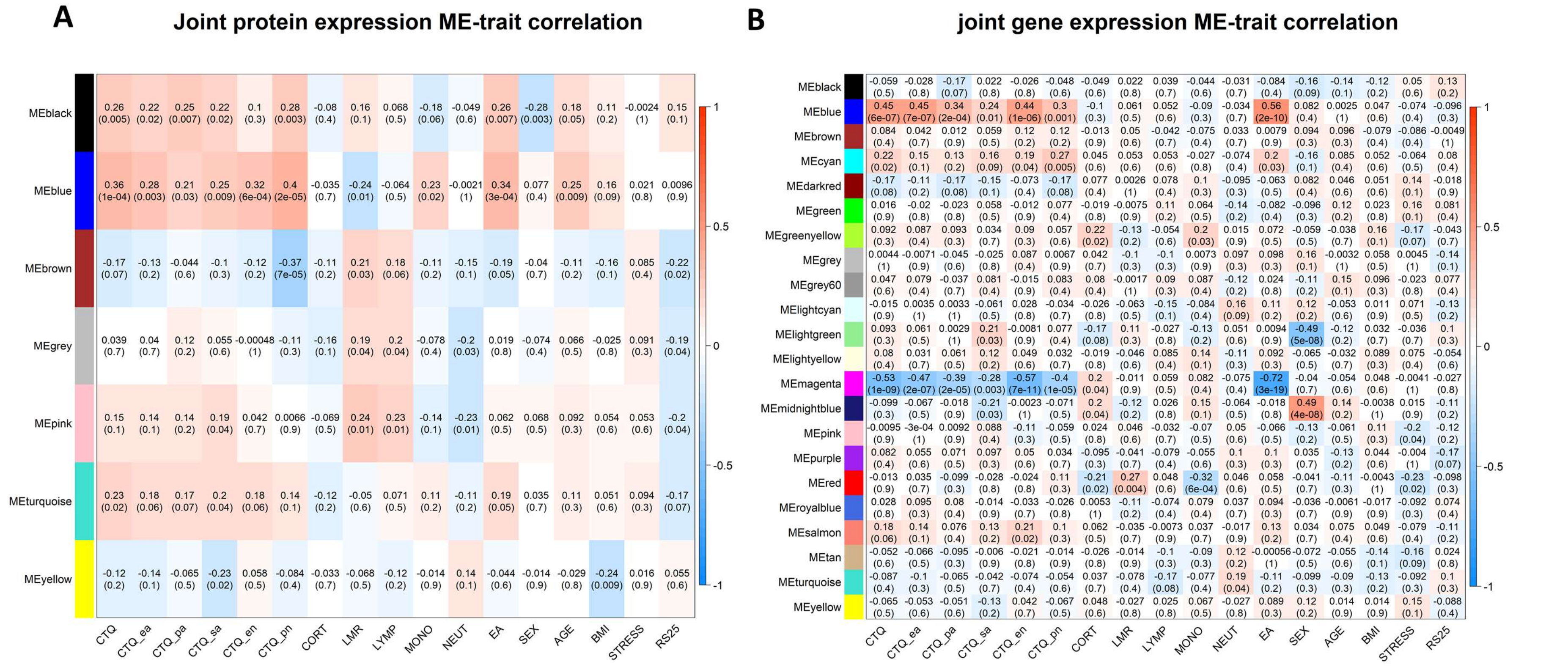


Figure S5. Module eigenproteins, eigengenes or eigenCpGs (ME) correlation with clinical variables are depicted for the joint protein (A) and gene expression (B) consensus network. Abbreviations: EA: early adversity, STRESS: baseline or post stress measurement, CTQ: Childhood Trauma Questionnaire total score (CTQ categories: sa = sexual abuse, pa = physical abuse, ea= emotional abuse, en = emotional neglect, pn = physical abuse), RS25: Resilience Scale 25 total score, CORT: Cortisol base to peak ratio, BMI: body mass index, SEX: biological sex, AGE: age in years at participation, LYMPH & MONO: count of lymphocytes and monocytes, LMR: lymphocyte to monocyte ratio.

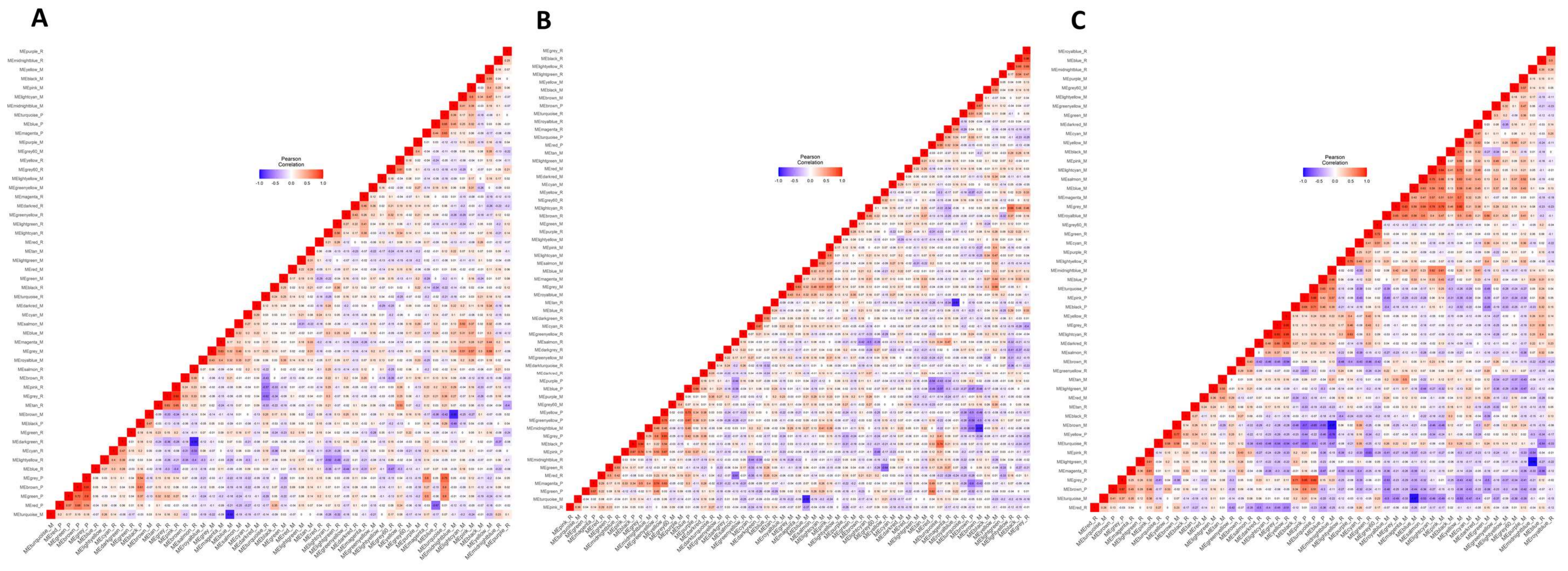


Figure S6. Heatmaps display clustered (HCA) correlation coefficients representing within and cross-omic interrelation of co-expression and co-methylation modules derived from baseleine (A) post stress (B) or joint analyses (C). Red indicates positive correlations, blue indicates negative correlations.

**Analyte specific co-expression and co-methylation structures and cross-omic interrelation**

Of all baseline modules, expression of the brown gene co-expression module ($r$ = .57, $p$ <.001 ) and the blue protein co-expression module ($r$ = 0.3, $p$ <.05) were significantly positively correlated with adversive childhood experience (Figure X). Further a magenta protein co-expression ($r$ = 0.26, $p$ = .05) and a green yellow gene co-expression module ($r$ = 0.25, $p$ = .05) were positively but not significantly associated with adversive childhood experience.

Transcripts contained within the brown baseline gene co-expression module were functionally implicated in e.g. DNA strand elongation (GO:0022616) or positive regulation of interleukin-1 beta production (GO:0032731), and green yellow gene co-expression module was enriched with transcripts playing a role in platelet degranulation (GO:0002576), chemokine-mediated signaling pathway (GO:0070098) regulation of macrophage derived foam cell differentiation (GO:0010743). The relationship between module expression and early adversity was comparatively lower in protein modules than in gene expression modules.

Baseline protein modules tended to show a lower correlation of expression and early adversity in comparison to gene co-expression modules. The blue protein co-expression module harbored proteins associated with e.g. neutrophil degranulation (GO: 0043312), viral process (GO:0016032), or mRNA processing (GO:0006397). The magenta protein co-expression module on the other side was enriched for proteins implicated in mitochondrial ATP synthesis coupled electron transport (GO:0042775) and the respiratory electron transport chain (GO:0022904) as well as viral gene expression (GO:0019080).

Following stress exposure, the number of adversity associated co-expression modules increased. Post stress gene co-expression modules (n = 5) were functionally associated with e.g. regulation of cellular response to heat (GO:1900034) and regulation of cellular response to stress (GO:0080135) (the salmon module, $r$ = 0.3, $p$ <.05), purine nucleoside monophosphate biosynthetic process (GO:0009127) and mitochondrial protein processing (GO:0034982) (the red module $r$ = 0.3, $p$ <.05), secretion (GO:0046903) (the light cyan module $r$ = 0.3, $p$ <.05), pyrimidine-containing compound transmembrane transport (GO:0072531) (the dark green module $r$ = 0.3, $p$ <.05) or acute inflammatory response (GO:0002526) (the brown module $r$ = 0.3, $p$ <.05).

Post stress protein co-expression modules (n = 3) were functionally associated with e.g. regulation of mRNA stability (GO:0043488) and antigen processing and presentation of exogenous peptide antigen via MHC class I, TAP-dependent (GO:0002479) (the turquoise module $r$ = -0.29, $p$ <.05), neutrophil degranulation (GO:0043312), neutrophil activation involved in immune response (GO:0002283) and viral release from host cell (GO:0019076) (the purple module $r$ = 0.35, $p$ <.01) or neutrophil degranulation (GO:0043312) and respiratory electron transport chain (GO:0022904) (the blue module $r$ = 0.35, $p$ <.01).

Notably, modules correlated with early adversity were often as well associated with participants' CTQ scores. None of the identified co-methylation modules was associated with the experience of childhood adversity and we found no relation between protein co-expression and stress experience. A red ($r$ = -0, 16, $p$ <.05) and a pink ($r$ = -0, 20, $p$ <.05) gene co-expression module however were slightly but significantly negatively associated with stress exposure. Transcripts of the red gene co-expression module are functionally invovled in e.g. the purine nucleoside monophosphate biosynthetic process (GO: 0009127) or mitochondrial protein processing (GO: 0034982). Transcripts contained in the pink module seemed to play a role in immune related processes and are functionally associated with T cell activation (GO: 0042110), B cell activation (GO: 0042113) or lymphocyte differentiation (GO: 0030098).

Correlational analysis related expression some of identified co-expression modules with the experience of childhood adversity. Yet, the experience of early adversity did not account for much of variance in module expression in particularly protein co-expression modules when accounting for the influence of other variables especially participants' CTQ scores. Multiple regression models, however, marked the salmon and the lightycan post stress gene expression module, the light yellow and brown baseline gene co-expression as well brown and the light yellow module identified in the joint gene expression analysis as potential modules of interest for the study of molecular effects of childhood adversity.

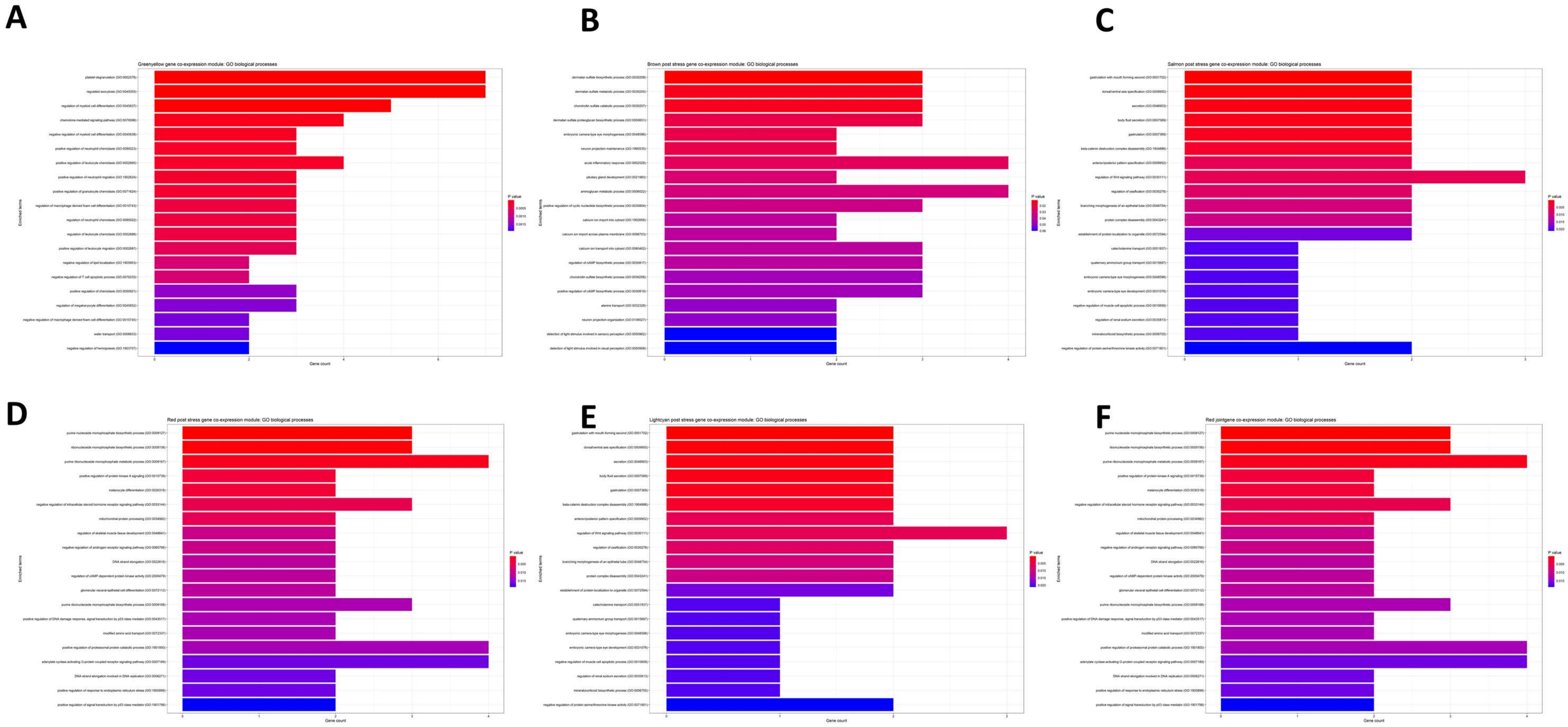


Figure S7. EnrichR derived biological processes as associated with analytes contained in gene co-expression modules of interest. Bar plots give number of analytes linked to the given GO-Term, colors represent levels of significance (red = smaller pvalues).Enrichment analyses are shown for the green yellow baseline module (A), the brown post stress module (B), the salmon post stress module (C), the red post stress module (D), the lightycan post stress module (E) and the red module derived from the joint gene co-expression analysis (F).

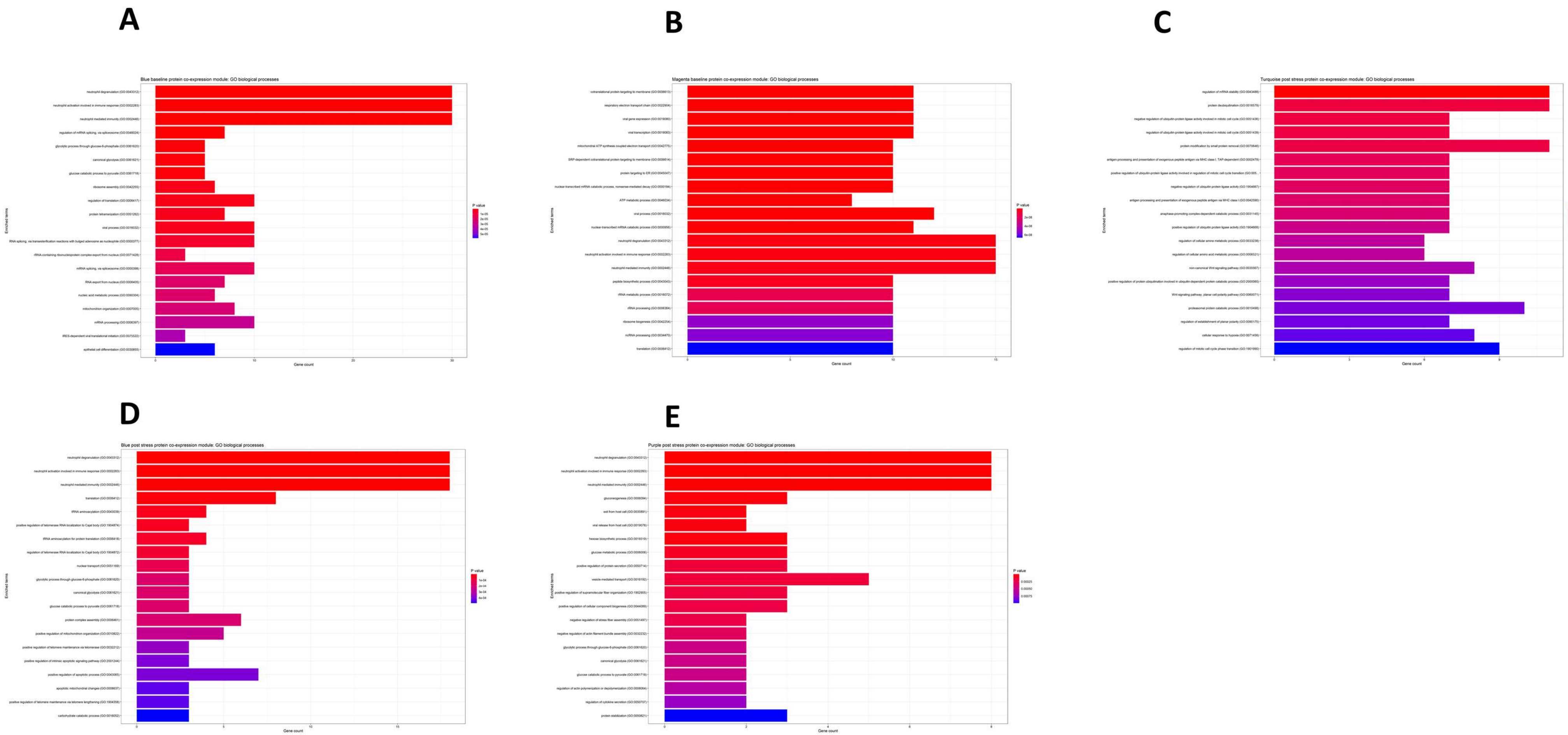


Figure S8. EnrichR derived biological processes as associated with analytes contained in protein co-expression modules of interest. Bar plots give number of analytes linked to the given GO-Term, colors represent levels of significance (red = smaller pvalues).Enrichment analyses are shown for the blue baseline module (A), the magma baseline module (B), the turquoise post stress module (C), the blue post stress module (D) and the purple post stress module.

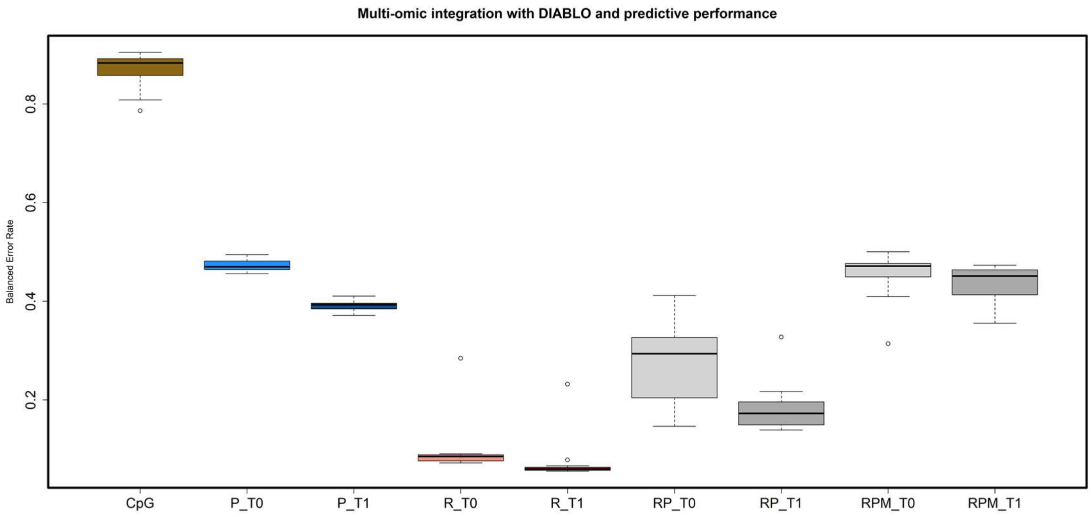


Figure S9 : We fit different models of varying complexity (10 models with 10 -100 increasing number of analytes selected) using DIABLOs block splsda (grey) to integrate transcriptomic and proteomic (RP) or transcriptomic, proteomic and CpG methylation data (RPM) on the individual datasets (yellow = CpG methylation (M), blue = protein expression data or red = gene expression data (R)) using sPLS Predictive performance was assessed through the balanced error rate. Post stress modules (T1) differentiated better between conditions (EA/CG) compared to baseline modules (T0) and we achieved the best performance with the sPLS model retaining 20 transcripts.

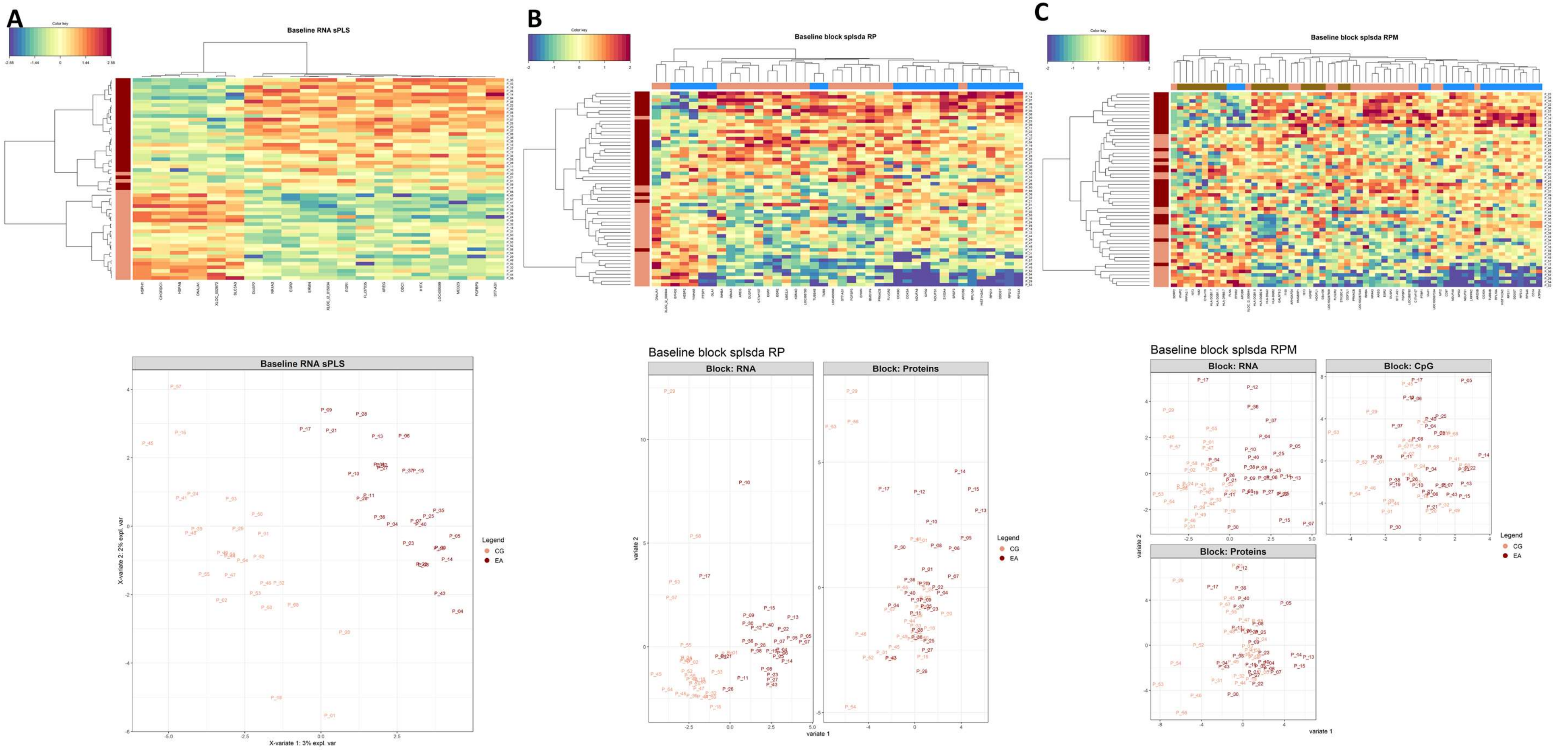


Figure S10. Top performing baseline regression (sPLS) and DIABLO models (block sPLS). The model with the best discriminant capabilities (A) is based on 20 transcripts derived from CD14⁺ monocytes isolated 180 min after stress exposure. Including post stress proteomic data (B) and CpG methylation data did not result in a variable selection that differentiates better between participants' with a history of childhood adversity and control participants. Heatmaps visualizes clustered analytes (columns) and participants (rows). Row annotations indicate participants' condition (EA = dark red). Scatterplots visualize participants' loadings (EA = dark red) on the first component extracted from analytes of included omic levels (A = RNA, B = RNA & Protein, C = RNA, Protein and CpG Methylation). Bar plots give biological processes associated with the analytes contained in the respective models. GO-Terms are derived from the Enrichr database.

Table S10: Transcripts included in the best performing supervised baseline gene-expression model

| **Transcripts** | | **Overall, N = 56** | **CG, N = 27** | **EA, N = 29** | **Difference1** | **95% CI1,2** | **p-value1** | **q-value3** |
|---|---|---|---|---|---|---|---|---|
| HSPH1 | Mean (SD) | 11.39 (0.40) | 11.67 (0.37) | 11.13 (0.21) | 0.54 | 0.38, 0.70 | <0.001 | <0.001 |
| CHORDC1 | Mean (SD) | 9.20 (0.28) | 9.37 (0.22) | 9.03 (0.21) | 0.34 | 0.22, 0.46 | <0.001 | <0.001 |
| HSPA8 | Mean (SD) | 13.21 (0.31) | 13.42 (0.23) | 13.02 (0.25) | 0.40 | 0.27, 0.53 | <0.001 | <0.001 |
| DNAJA1 | Mean (SD) | 12.34 (0.28) | 12.55 (0.22) | 12.15 (0.17) | 0.41 | 0.30, 0.51 | <0.001 | <0.001 |
| XLOC_002872 | Mean (SD) | 8.31 (0.26) | 8.47 (0.23) | 8.16 (0.19) | 0.32 | 0.20, 0.43 | <0.001 | <0.001 |
| SLC5A3 | Mean (SD) | 9.86 (0.34) | 10.07 (0.25) | 9.67 (0.30) | 0.40 | 0.26, 0.55 | <0.001 | <0.001 |
| DUSP2 | Mean (SD) | 6.83 (0.53) | 6.50 (0.36) | 7.14 (0.47) | -0.64 | -0.86, -0.42 | <0.001 | <0.001 |
| NR4A3 | Mean (SD) | 4.04 (1.53) | 2.94 (0.78) | 5.08 (1.32) | -2.1 | -2.7, -1.6 | <0.001 | <0.001 |
| EGR2 | Mean (SD) | 7.49 (0.83) | 6.88 (0.59) | 8.06 (0.56) | -1.2 | -1.5, -0.87 | <0.001 | <0.001 |
| ERMN | Mean (SD) | 5.64 (0.47) | 5.30 (0.36) | 5.95 (0.34) | -0.65 | -0.84, -0.46 | <0.001 | <0.001 |
| XLOC_l2_015034 | Mean (SD) | 6.42 (0.59) | 6.00 (0.43) | 6.82 (0.41) | -0.82 | -1.1, -0.60 | <0.001 | <0.001 |
| EGR1 | Mean (SD) | 10.28 (0.91) | 9.64 (0.79) | 10.88 (0.54) | -1.2 | -1.6, -0.87 | <0.001 | <0.001 |
| FLJ37035 | Mean (SD) | 7.28 (0.27) | 7.10 (0.22) | 7.44 (0.19) | -0.34 | -0.45, -0.23 | <0.001 | <0.001 |
| AREG | Mean (SD) | 4.50 (1.63) | 3.22 (0.65) | 5.70 (1.33) | -2.5 | -3.0, -1.9 | <0.001 | <0.001 |
| ODC1 | Mean (SD) | 11.29 (0.44) | 11.00 (0.38) | 11.56 (0.30) | -0.56 | -0.74, -0.37 | <0.001 | <0.001 |
| H1FX | Mean (SD) | 11.04 (0.33) | 10.82 (0.25) | 11.23 (0.26) | -0.41 | -0.55, -0.27 | <0.001 | <0.001 |
| LOC400099 | Mean (SD) | 10.66 (0.28) | 10.46 (0.21) | 10.84 (0.20) | -0.37 | -0.49, -0.26 | <0.001 | <0.001 |
| MED23 | Mean (SD) | 7.56 (0.30) | 7.36 (0.23) | 7.74 (0.24) | -0.37 | -0.50, -0.25 | <0.001 | <0.001 |
| FGFBP3 | Mean (SD) | 5.90 (0.33) | 5.67 (0.27) | 6.12 (0.19) | -0.46 | -0.58, -0.33 | <0.001 | <0.001 |
| ST7-AS1 | Mean (SD) | 4.89 (0.61) | 4.48 (0.45) | 5.28 (0.48) | -0.80 | -1.0, -0.55 | <0.001 | <0.001 |

1 Welch Two Sample t-test

2 CI = Confidence Interval

3 False discovery rate correction for multiple testing

### References


Dieckmann, L., Cole, S., & Kumsta, R. (2020). Stress genomics revisited: Gene co-expression analysis identifies molecular signatures associated with childhood adversity. *Translational Psychiatry*, *10*(1), 34. https://doi.org/10.1038/s41398-020-0730-0

Epskamp, S., Cramer, A. O. J., Waldorp, L. J., Schmittmann, V. D., & Borsboom, D. (2012). qgraph : Network Visualizations of Relationships in Psychometric Data. *Journal of Statistical Software*, *48*(4). https://doi.org/10.18637/jss.v048.i04

Frach, L., Tierling, S., Schwaiger, M., Moser, D., Heinrichs, M., Hengstler, J., . . . Kumsta, R. (2019). *The mediating role of KITLG DNA methylation in the association between childhood adversity and cortisol stress reactivity does not replicate in monocytes.* https://doi.org/10.31234/osf.io/6ed7n

Franceschini, A., Szklarczyk, D., Frankild, S., Kuhn, M., Simonovic, M., Roth, A., . . . Jensen, L. J. (2012). String v9.1: Protein-protein interaction networks, with increased coverage and integration. *Nucleic Acids Research*, *41*(Database issue), D808-15. https://doi.org/10.1093/nar/gks1094

Langfelder, P., & Horvath, S. (2008). Wgcna: An R package for weighted correlation network analysis. *BMC Bioinformatics*, *9*, 559. https://doi.org/10.1186/1471-2105-9-559

Langfelder, P., Luo, R., Oldham, M. C., & Horvath, S. (2011). Is my network module preserved and reproducible? *PLoS Computational Biology*, *7*(1). https://doi.org/10.1371/journal.pcbi.1001057

Rohart, F., Gautier, B., Singh, A., & Lê Cao, K.-A. (2017). Mixomics: An R package for 'omics feature selection and multiple data integration. *PLoS Computational Biology*, *13*(11), e1005752. https://doi.org/10.1371/journal.pcbi.1005752

Schwaiger, M., Grinberg, M., Moser, D., Zang, J. C. S., Heinrichs, M., Hengstler, J. G., . . . Kumsta, R. (2016). Altered stress-induced regulation of genes in monocytes in adults with a history of childhood adversity. *Neuropsychopharmacology : Official Publication of the American College of Neuropsychopharmacology*, *41*(10), 2530–2540. https://doi.org/10.1038/npp.2016.57

Speicher, N. K., & Pfeifer, N. (2015). Integrating different data types by regularized unsupervised multiple kernel learning with application to cancer subtype discovery. *Bioinformatics (Oxford, England)*, *31*(12), i268-75. https://doi.org/10.1093/bioinformatics/btv244

Stelzer, G., Rosen, N., Plaschkes, I., Zimmerman, S., Twik, M., Fishilevich, S., . . . Lancet, D. (2016). The GeneCards Suite: From Gene Data Mining to Disease Genome Sequence Analyses. *Current Protocols in Bioinformatics*, *54*, 1.30.1-1.30.33. https://doi.org/10.1002/cpbi.5

Zang, J.C.S.*, May, C.*, Hellwig, B., Moser, D., Hengstler, J.G., Cole, S., . . . Kumsta, R. Proteome analysis of monocytes implicates altered mitochondrial functioning in adults reporting adverse childhood experiences. *Manuscript Submitted*, *2021*.
